# Modulation of SARS-CoV-2 spike binding to ACE2 through conformational selection

**DOI:** 10.1101/2024.03.15.585207

**Authors:** Prithwidip Saha, Ignacio Fernandez, Fidan Sumbul, Claire Valotteau, Dorota Kostrz, Annalisa Meola, Eduard Baquero, Arvind Sharma, James R. Portman, François Stransky, Thomas Boudier, Pablo Guardado Calvo, Charlie Gosse, Terence Strick, Felix A. Rey, Felix Rico

## Abstract

The first step of SARS-CoV-2 infection involves the interaction between the trimeric viral spike protein (*S*) and the host angiotensin-converting enzyme 2 (*ACE*2). The receptor binding domain (*RBD*) of *S* adopts two conformations: open and closed, respectively, accessible and inaccessible to *ACE*2. Therefore, *RBD* motions are suspected to affect *ACE*2 binding; yet a quantitative description of the underlying mechanism has been elusive. Here, using single-molecule approaches, we visualize *RBD* opening and closing and probe the *S*/*ACE*2 interaction. Our results show that *RBD* dynamics affect *ACE*2 binding but not unbinding. The resulting modulation is quantitatively predicted by a conformational selection model in which each protomer behaves independently. Our work reveals a general molecular mechanism affecting binding affinity without altering binding strength, helping to understand coronavirus infection and immune evasion.

## Main text

The first step in the entry of SARS-CoV-2 into a target cell is recognition by the viral spike protein (*S*) of the human angiotensin-converting enzyme 2 (*ACE*2) at the plasma membrane of target cells. The *S* protein forms a trimer that undergoes maturation to yield two subunits, *S*1 (forming the head of the spike and responsible for recognition) and *S*2 (forming the stalk and responsible for subsequent membrane fusion) (*1, 2*). The receptor binding domain (*RBD*) is part of *S*1 and has been shown to adopt two distinct conformations, open and closed (*3*–*9*). Only the open *RBD* interacts and forms a complex with *ACE*2. This interaction has been probed using both ensemble and single-molecule binding assays, showing that the isolated *RBD* displays higher affinity for *ACE*2 compared with the *RBD* embedded within the *S* trimer (*3, 10*–*17*). *RBD* switching between open and closed conformations explain this observation (*18*). Supporting this idea, it has been shown that the occurrence of *S* variants with mutations away from the receptor binding motif modulates binding to *ACE*2 and binding is suppressed when disulfide bonds are introduced to lock *RBD* in the closed conformation (*19*–*24*). Microsecond-scale molecular dynamics simulations revealed large conformational changes in *S*1 and elucidated the impact of glycans and *RBD*-distal mutations in the closed/open transition pathways (*25*–*28*). Experimentally, single-molecule FRET (smFRET) monitored real-time dynamics of labeled *RBD* in *S* trimers, revealing subsecond transition times between up to four conformations attributed to states in which one or multiple *RBD*s are open (*29*–*32*). Finally, high-speed atomic force microscopy (HS-AFM) imaging monitored the dynamics of these transitions (*17, 33, 34*). Although no rate constants were reported, it clearly showcased the potential of this technique. Despite these important advances, various outstanding questions remain, including whether each *RBD* in the trimeric spike opens and closes independently or in coordination with the others, and how the thermodynamics and kinetics of the conformational change affect the affinity of *S* trimers for *ACE*2.

Here, we used HS-AFM imaging to visualize in real-time the conformational dynamics of all three *RBD*s of an *S* trimer, thereby measuring their opening and closing rate constants. We also proved that the *RBD* of each *S* protomer opens and closes independently. We then probed the bond strength of *ACE*2 to both the isolated *RBD* and the full-length *S* trimer over a wide range of conditions using complementary approaches: bilayer interferometry (BLI), magnetic tweezers (MT) coupled to junctured-DNA, and HS-AFM-based single-molecule force spectroscopy (HS-FS). All our results consistently showed that the parameters of the energy landscape towards dissociation and the corresponding rate constant at zero force are similar for an isolated *RBD* and for a *RBD* within the *S* trimer. Conversely, we found that the binding of *ACE*2 to *S* trimers is modulated by the fast *RBD* conformational changes, which affect both the association rate constant and the affinity. Our data derived from multiple experimental techniques are quantitatively and coherently described by a conformational selection model.

### Real-time conformational dynamics of *RBD*s

To quantify *RBD* opening and closing, we imaged *S* trimers on mica surfaces with HS-AFM, at subsecond timescale, nanometer resolution, and in near-physiological conditions (see Materials and Methods) (*35*). To improve the stability of recombinant *S* trimers, we used two engineered versions of the Wuhan variant, one with mutations at the furin protease cleavage site located between *S*1 and 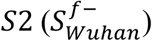, and the second with a double proline substitution in the loop between the first heptad repeat and the central helix of 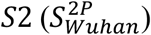, as encoded in various mRNA-based vaccines (*2, 3, 5, 36*). HS-AFM videos revealed up to three protrusions of 3-5-nm size, stochastically projecting out from the bulbous head of *S* trimers (Fig. 1A, figs. S2-S3 and supplementary videos). We interpreted them as the signature of *RBD* opening on each *S* protomer, in line with the simulated topographical images derived from cryo-EM structures (*37*) (Fig. 1A and see Materials and Methods). Thus, four *S*_*i*_ states were identified for the *S* trimer, with *i* = 0 to 3 indicating the number of open *RBD*s. We developed a semi-automatic image analysis routine to process thousands of images (see Materials and Methods and fig. S1), assigning each image to one of four identified states and yielding opening/closing trajectories over time (Fig. 1B).

**Fig. 1.**
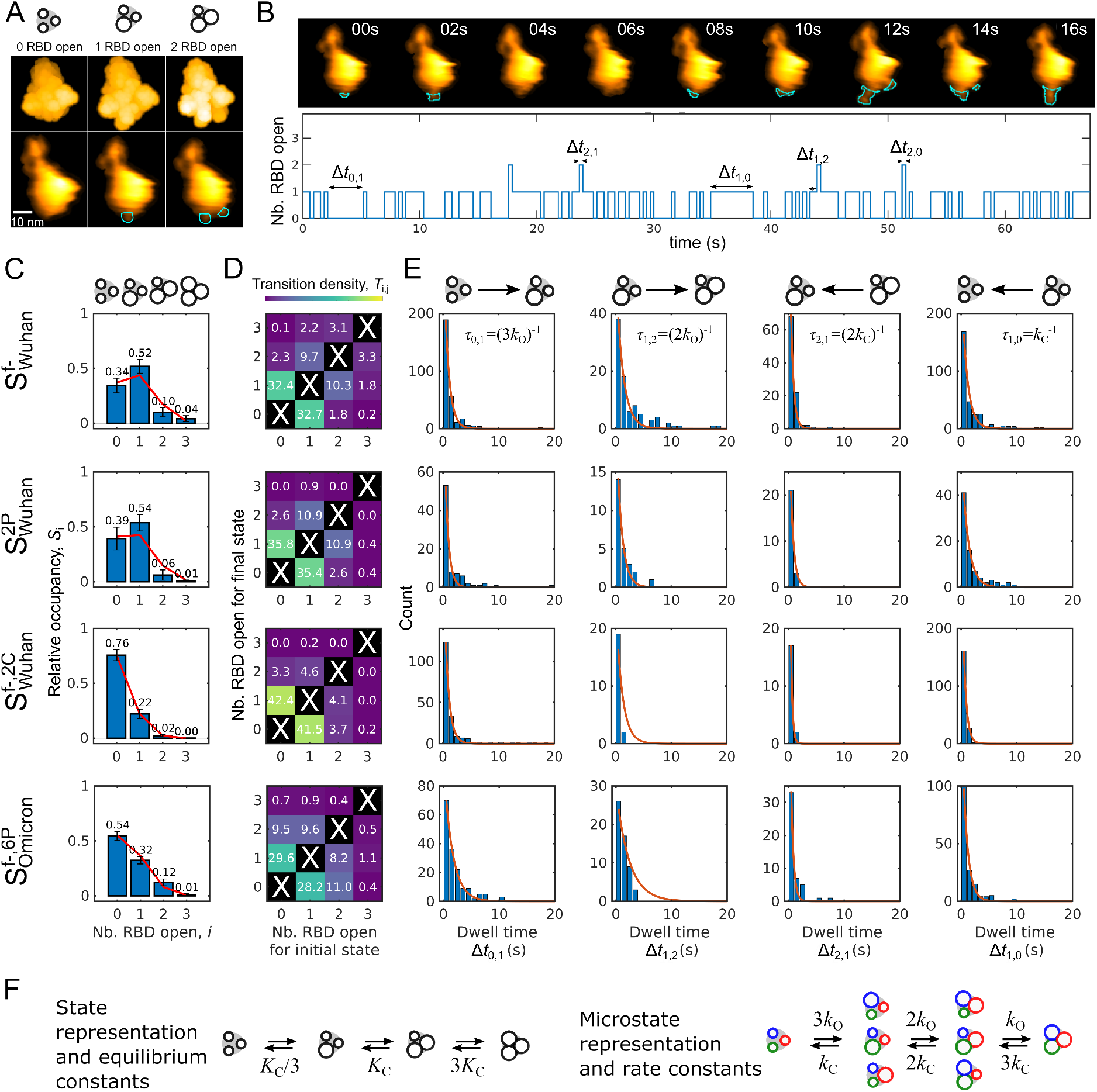
Single-molecule real-time visualization of the conformational dynamics of *RBD* opening and closing in *S* trimers. **(A)** Top. Simulated HS-AFM topography of an 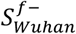 trimer immobilized on a mica surface showing 0 to 2 *RBD*s open (left to right, PDB: 7DDD, 7BNN, and 7BNO, respectively). Bottom. Corresponding experimental images. **(B)** Top. Representative frames extracted every 2 s from an HS-AFM video showing the opening and closing *RBD*s (cyan lines). Bottom. Representative trajectory showing the number of *RBD*s in the open state for one HS-AFM video acquired at 1 fps. **(C)** Relative occupancy histograms (mean ± SEM, blue bars) and associated fits to the binomial distribution (red lines) for the four *S*_*i*_ states, *i* indicating the number of opened *RBD*s. **(D)** Transition density plots (TDP) showing the fraction of transition events between the different states. **(E)** Distributions of the Δ*t*_*i,j*_ dwell times for the four most populated *S*_*i*_ to *S*_*j*_ transitions. Red lines show global fits of exponential decays (see Table 1 for results and table S2 for statistics). **(F)** Schematic representation of all possible states and microstates for *S* trimers as well as of all possible transitions. *K*_*C*_ is the *RBD* closing equilibrium constant, and *k*_*O*_ and *k*_*C*_ are the corresponding opening and closing rate constants, respectively. The *RBD* coloration for the microstates is pedagogical, the protomers being indistinguishable.

From these trajectories, we then computed the relative occupancy of each state *S*_*i*_ and displayed them as state density histograms (Fig. 1C). The results for 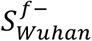 and 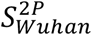 revealed that the *S*_1_ state was the most populated, with more than half of the total (52 and 54 %, respectively), followed by the *S*_0_ (34 and 39 %) and *S*_2_ states (10 and 6 %), the *S*_3_ state being the least populated one (<5 %). Particle classification of Wuhan variant *S* trimer structures from cryo-EM agrees with our results (supplementary text S1) (*2, 5, 7*). We obtained rough estimates of the *RBD* closing equilibrium constants, *K*_*C*_, by first computing the ratios between the populations of closed and open *RBD*s (Table 1), yielding 2.6 ± 0.6 for 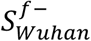 and 3.3 ± 0.8 for 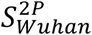. Next, to test the assumption of independent opening between the different protomers, we fitted the state density histograms with a binomial distribution and obtained the probabilities for an *RBD* to be open (*p*_*open*_) (Fig. 1C, Table 1). These measurements confirmed our hypothesis of independence of the transitions. Alternatively, computing *K*_*C*_ as *p*_*closed*_ /*p*_*open*_ = 1/*p*_*open*_ − 1 led to 2.5 ± 0.4 and 2.9 ± 0.6 for 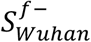 and 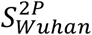, respectively, in agreement with our rough estimates.

**Table 1.** Kinetic and thermodynamic parameters for the opening and closing transitions of an individual *RBD* within the *S* trimer. The closing equilibrium constant, *K*_*C*_, was calculated from three different approaches. First, from the ratio of total number of *RBDs* observed in the closed conformation (Σ*RBD*_*closed*_ = Σ_*i*_ (3 − *i*) × *S*_*i*_) over the total number of *RBDs* observed in the open state (Σ*RBD*_*open*_ = ∑_*i*_ *i* × *S*_*i*_). Second, from the probability for an *RBD* to be in the open state, as obtained from the fit of the binomial distribution 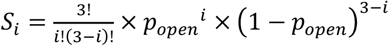 to the relative occupancy histograms (Fig. 1C). Third, from the ratio between the closing and opening rates extracted from the global fits to the dwell time distributions (Fig. 1E).

| *S*<br>variant | $K_C \equiv \frac{\Sigma RBD_{closed}}{\Sigma RBD_{open}}$ | $p_{open}$ | $K_C \equiv \frac{1}{p_{open}} - 1$ | $k_C$ (s<sup>-1</sup>) | $k_O$ (s<sup>-1</sup>) | $K_C \equiv \frac{k_C}{k_O}$ |
| --- | --- | --- | --- | --- | --- | --- |
| $S_{Wuhan}^{f-}$ | $2.6 \pm 0.6$ | $0.28 \pm 0.04$ | $2.5 \pm 0.4$ | $1.00 \pm 0.28$ | $0.40 \pm 0.11$ | $2.5 \pm 1.4$ |
| $S_{Wuhan}^{2P}$ | $3.3 \pm 0.8$ | $0.26 \pm 0.04$ | $2.9 \pm 0.6$ | $0.88 \pm 0.36$ | $0.52 \pm 0.21$ | $1.7 \pm 1.4$ |
| $S_{Wuhan}^{f-,2C}$ | $10.5 \pm 2.0$ | $0.09 \pm 0.00$ | $10.4 \pm 0.0$ | $1.81 \pm 0.60$ | $0.44 \pm 0.15$ | $4.2 \pm 2.8$ |
| $S_{Omicron}^{f-,6P}$ | $4.0 \pm 0.6$ | $0.18 \pm 0.01$ | $4.6 \pm 0.4$ | $1.12 \pm 0.26$ | $0.21 \pm 0.05$ | $5.2 \pm 2.5$ |

HS-AFM videos also provided access to *RBD* opening/closing dynamics. From the pooled opening/closing trajectories (Fig. 1B), we calculated the probability of transitions from state *S*_*i*_ to state *S*_6_ to build transition density plots (TDP, Fig. 1D) (*38*). The data displayed the diagonal symmetry expected from the detailed balance between conformational transitions at equilibrium. The higher frequencies of transitions for |*j* − *i*| = 1 (>85 %) are easy to understand given the relative occupancy of the various states. We attributed the uncommon |*j* − *i*| > 1 transitions to the limited time resolution of HS-AFM.

To get quantitative insight into opening and closing kinetics, we generated histograms of the Δ*t*_*i,j*_ dwell times before the four most populated *S*_*i*_ to *S*_*j*_ transitions (representing 85 – 95 % of the total, Fig. 1E). Exponential fitting of the individual histograms yielded *τ*_*i,j*_ characteristic times of ∼2 s before opening, and ∼3 s before closing. Assuming independence between protomers, the observed transition rate constants, *k*_*ij*_ = 1/*τ*_*i,j*_, are related to the opening and closing rates of single *RBD, k*_*O*_ and *k*_*C*_, by *k*_*i,i*+1_ = (3 − *i*) × *k*_*O*_ and *k*_*i*+1,*i*_ = (*i* + 1) × *k*_*C*_ (Fig. 1F). Thus, we globally fitted the 4 histograms with monoexponential decays with the opening and closing rate constants and four transition amplitudes as free parameters. This yielded opening rates about twice slower than closing rates *(k*_*O*_ *=* 0.4 – 0.5 s^-1^ and *k*_*C*_ = 0.9 – 1 s^-1^) for 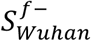 and 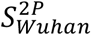 (Table 1), *i*.*e*. the time of an *S* protomer with its *RBD* open is about half of that with the *RBD* closed. Recast in terms of closing equilibrium constants *K*_*C*_ = *k*_*C*_/*k*_*O*_, we obtained 2.5 ± 1.4 for 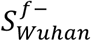 and 1.7 ± 1.4 for 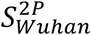 (Table 1), in agreement with population estimates. Overall, from both thermodynamic and kinetic points of view, the two constructs for the Wuhan spikes we studied do not differ from each other. The similarity of the results of the *f*– and 2*P* forms, despite displaying mutations at very different locations along the sequence, suggests that they do not affect the opening and closing of the *RBD*. Our approach allows quantification of effective opening and closing rates for each individual *RBD* in the *S* trimer and suggests conformational changes independent of each other. Population and kinetic analysis lead to similar results and provide a framework that can be applied to other techniques, either from snapshots obtained by cryo-EM single particle analysis (table S1), or transition rates derived from smFRET.

To validate our HS-AFM approach, we imaged the conformational states of construct 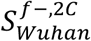, which has D985 and S383 replaced by cysteines, engineered to form a disulfide bond to retain the *RBD*s in the closed state, thereby impairing *ACE*2 binding (20, 39, 40). Compared to 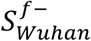, *RBD* closed states are favored: 76 ± 7 % of all spikes showed no open *RBD*s, 22 ± 4 % displayed one open *RBD*, and 2 ± 1 % displayed two open *RBD*s (Fig. 1C). A similar mutant also presenting the 2*P* mutation 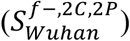 provided similar results, further confirming our approach and the minor effect of the 2*P* mutation (fig. S4, table S2). Although considerably reduced, some degree of *RBD* dynamics is still allowed after the 2*C* substitution, suggesting that there is not 100 % disulfide bond formation in this construct, which is consistent with cryo-EM studies (24, 40). Furthermore, while the opening rate remained unchanged, the closing rate was almost twice faster for 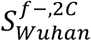 than that of 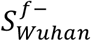 (Table 1). The faster rate closing constants of 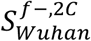 are similar to those found in smFRET experiments, suggesting possible intermediate states, not considered in our analysis (supplementary text S1) (29, 32). The probability of an *RBD* to be in the open state in the 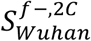 construct was 0.1 and *K*_*C*_ was estimated to be between 5 and 11 (Table 1). While the *K*_*C*_ values obtained from population and kinetic analyses were similar, their accuracy was limited by the frame rate of our HS-AFM imaging (1-2.5 frame per second), which was similar to the closing rate constant. Therefore, for 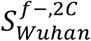, the population analysis might be more accurate.

One of the most prevalent variants of concern of SARS-CoV-2 is Omicron (B.1.1.529), which presents over 30 amino acid mutations compared to the Wuhan strain (*23, 41*). We used the structure-based designed HexaPro construct of the Omicron spike (*19*). The 6 prolines introduced in this construct allow a much higher yield of recombination. The relative occupancy derived from HS-AFM imaging showed important differences compared to Wuhan 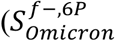, fig. S5, and supplementary videos), with over 54 % of the population in the all *RBD* closed state, and ∼32 % with one *RBD* open (Fig. 1C). While cryo-EM studies have reported contradictory results, several studies agree with ours (table S1) (*22, 40, 42*–*44*). The TDPs also revealed a second distinction between the Wuhan and Omicron spikes, with transitions involving more than one *RBD* being more frequent in 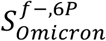 (∼25 %) than in 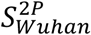 (<9 %) (Fig. 1D). While this result may suggest some cooperativity in the Omicron *RBD* opening and closing, the state density histogram was still perfectly fitted with a binomial distribution, which speaks up for independent conformational changes. Additionally, global fit analysis of dwell time histograms revealed an opening rate approximately twice slower than for Wuhan spike and a slightly faster closing rate (Table 1). Our results in the Wuhan variants suggested no significant differences in *RBD* dynamics due to the 2*P* substitution. Thus, the dynamics of the Omicron *S* trimers were likely not affected by the 6*P* substitution, in line with the preserved immunogenicity of this construct (*19*). Therefore, our observed difference in *RBD* state occupancy and dynamics between Omicron and Wuhan variants is likely due to the acquired mutations during SARS-CoV-2 evolution in the human population, as recently suggested by molecular dynamics simulations (*45*). Moreover, the higher population of closed states in Omicron would explain the lower multivalent binding to *ACE*2, as recently described (*17*).

### Binding strength and affinity to *ACE*2

To quantify the possible effect of opening and closing of *RBD* on its binding to *ACE*2, we compared the complexation of *ACE*2 to isolated *RBD* and to the 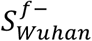 and 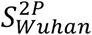 trimers using biolayer interferometry (BLI) (Fig. 2). We analyzed the time traces using a simple 1:1 binding model. The results showed similar dissociation rate constants of 3-4 × 10^-3^ s^-1^ (Table 2) for both an isolated *RBD* and a *RBD* in an *S* trimer. Conversely, *ACE*2 showed a ∼3-fold lower association rate constant for *RBD* within 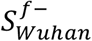 and 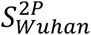 than for isolated *RBD* (Table 2, note that in the former case, we consider *apparent* association rate constants because the conformational change kinetics may modulate association). Therefore, *RBDs* within *S* trimers showed a 3-fold higher apparent equilibrium dissociation constant 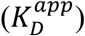 than isolated *RBDs* (Table 2).

**Table 2.** Kinetic and thermodynamic parameters for *ACE2* binding to the *RBD*, either isolated or embedded within the *S* trimer. *k*_*D*_ and *k*_*A*_ respectively correspond to the dissociation and association rate constants, *K*_*D*_ to the *k*_*D*_ /*k*_*A*_ dissociation equilibrium constant. Whereas all three parameters refer to microscopic events in the *RBD* case, they refer to a phenomenological 1:1 model in the *S* trimer (“app” = apparent). *K*_*C*_ is the *RBD* closing equilibrium constant that can be deduced from the measured dissociation equilibrium constants thanks to the conformational selection model (see below). BLI experiments (Fig. 2) were performed at room temperature in PBS; MT experiments at constant force (Fig. 4) were performed at 30 °C in 10 mM Na-HEPES pH 7.4, 150 mM NaCl, 0.1 % Tween 20, and 0.5 mg/mL BSA.

| Setup | Viral partner | $k_D$<br>($10^{-3} \text{ s}^{-1}$) | $k_A$<br>(or $k_A^{app}$ for *S*)<br>($10^5 \text{ M}^{-1}\text{s}^{-1}$) | $K_D$<br>(or $K_D^{app}$ for *S*)<br>(nM) | $K_C \equiv \frac{K_D^{app}}{K_D} - 1$ |
| --- | --- | --- | --- | --- | --- |
| BLI | *RBD*<sub>Wuhan</sub> | $3.0 \pm 0.0$ | $3.8 \pm 0.0$ | $7.8 \pm 0.1$ | |
| | *S*<sub>Wuhan</sub><sup>2P</sup> | $4.3 \pm 0.0$ | $1.7 \pm 0.0$ | $24.5 \pm 0.3$ | $2.1 \pm 0.1$ |
| | *S*<sub>Wuhan</sub><sup>f-</sup> | $4.1 \pm 0.0$ | $1.5 \pm 0.0$ | $26.9 \pm 0.3$ | $2.5 \pm 0.1$ |
| MT | *RBD*<sub>Wuhan</sub> | $21.3 \pm 1.4$ | $9.0 \pm 1.5$ | $23.7 \pm 3.6$ | |
| | *S*<sub>Wuhan</sub><sup>2P</sup> | $30.1 \pm 2.3$ | $3.6 \pm 0.5$ | $83.5 \pm 10.7$ | $2.5 \pm 0.5$ |

**Fig. 2.**
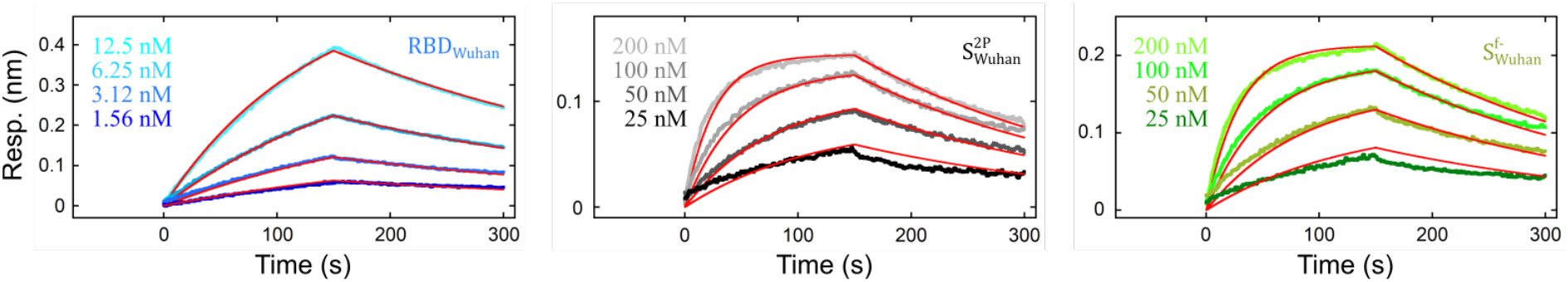
Ensemble measurements of the kinetic and thermodynamic parameters for *ACE2* binding to the *RBD*, either isolated or embedded within the *S* trimer. Bilayer interferometry response signals were obtained after injection of *ACE*2 at the indicated concentrations (from 0 to 150 s) and subsequent washing with buffer (from 150 to 300 s) for surfaces coated with either *RBD*_*Wuhan*_ (blue to cyan), 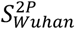 (black to grey), or 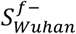 (green to lime). Red lines were fitted using a simple 1:1 binding model (see Table 2 for results).

BLI measurements were made with minor mechanical stress applied to the analytes. However, during normal breathing and even more during pathological coughing, the airflow in the respiratory tract applies important viscous drag forces to virus particles, affecting the interaction of the *S* trimers with *ACE*2 (*3, 10, 14, 23*). Recent single-molecule force spectroscopy (SMFS) probed the forces required to rupture *RBD* and *S* trimer/*ACE*2 complexes, proposing force-tuned avidity (*17*). Various SMFS reports have compared for different SARS-CoV-2 variants the influence of force on the stability of the complexes formed by *ACE*2 with both the *RBD* or an *S* trimer (*11, 12, 15*). However, they have only explored a restricted range of loading rates whereas under physiological conditions, the shear rate applied by the airflow varies over 7 orders of magnitude (*46*). Hence, we conducted HS-FS to measure the rupture forces of *RBD* and *S* trimers in complex with *ACE*2, over 5 decades of force loading rates (Fig. 3A-C) (*47, 48*). To covalently immobilize the molecules to the surfaces, we engineered *RBD, S* trimers, and *ACE*2 with a C-terminal ybbR tag (Fig. 3A) (*49*). This strategy enables site-directed grafting through a 32-nm polyethylene glycol flexible linker that allows the molecules to freely align with the pulling vectors. Isolated *RBD* and *S* trimers unbinding from *ACE*2 showed similar rupture forces over the investigated range (Fig. 3C). The most probable rupture forces first increased linearly with the logarithm of the loading rate and showed an upturn at the highest values (Fig. 3C, see Materials and Methods). The free energy landscape parameters obtained from fitting the force spectra with the Cossio-Hummer-Szabo (CHS) model were similar: intrinsic dissociation rate *k*_*D*_∼10^-2^ s^-1^, distance to transition state *x*^‡^∼0.9 nm, and barrier height Δ*G*^‡^∼15 k_B_T (Fig. 3C, table S3) (*50*). This observation confirmed that the binding strength and dissociation rates are parameters characteristic of the *RBD* and are not importantly affected by their inclusion within the *S* trimer.

**Fig. 3.**
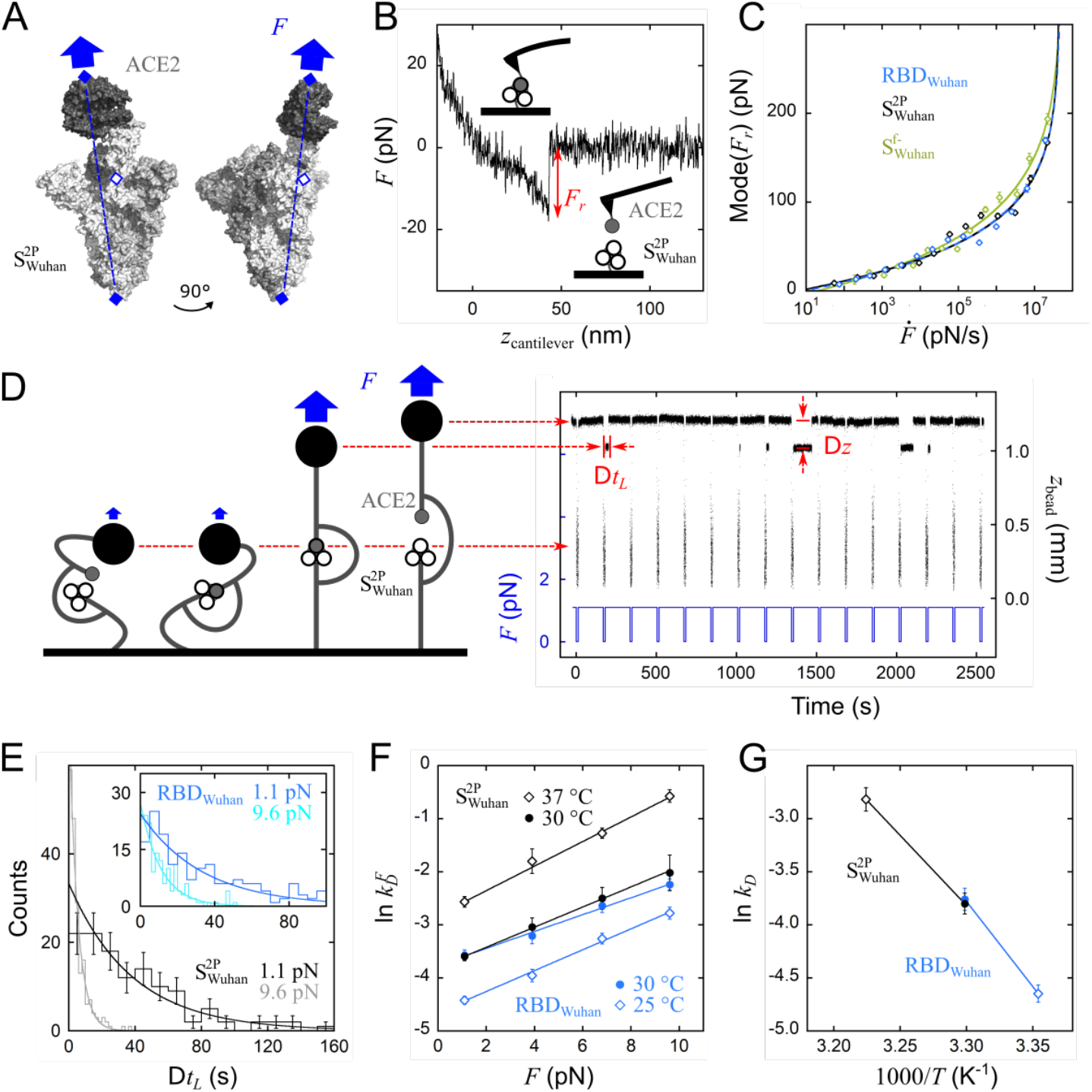
Single-molecule measurements of the dissociation kinetics and force response for *ACE2* bound to the *RBD*, either isolated or embedded within the *S* trimer. **(A)** Cryo-EM structure of the complex 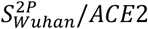 with one *RBD* open and bound (PDB 7A94 (*7*) visualized with PyMOL). The pulling direction (blue dashed line) connects the positions of the ybbR tags (full diamonds) used to tether *S* and *ACE*2. The open diamond is the position of the ybbR tag when pulling on isolated *RBD*. **(B)** Example of HS-AFM force-distance curve recorded on a 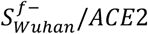 complex and showing how the rupture force, *F*_*r*_, is determined. **(C)** Dynamic force spectra from HS-AFM for *ACE*2 unbinding from *RBD*_*Wuhan*_, 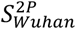, and 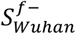 (blue, black, and kaki, respectively). The most probable rupture force was determined as the mode of the *F*_*r*_ distribution at each loading rate, 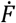 (figs. S6-S8; error bars are SEM). Fits to the CHS model (Eqs.1-2 in Materials and Methods) provided the position of the transition states in energy and space, Δ*G*^‡^ and *x*^‡^, as well as the dissociation rate constants at zero force, *k*_*D*_ (table S3). **(D)** Force-cycling procedure applied with MT to 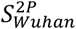 and *ACE*2 engrafted on J-DNA. At near 0 pN, the scaffold extension is low (two leftmost schemes, not to scale); the scaffold tips can encounter each other, and the two proteins can associate. When the force is increased to a value *F*, the bead pulls on the looped J-DNA and the extension reaches a higher value (middle scheme). Eventually, the two partners spontaneously separate and the extension increases again (right scheme). Upon repeating the force variation multiple times, one obtains a time-trace on which dissociation events are characterized by their dwell time in the looped conformation, Δ*t*_*L*_, and the amplitude of their extension jump, Δ*z*. **(E)** Examples of Δ*t*_*L*_ histograms collected on 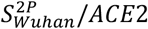 at *F* = 1.1 and 9.6 pN, *T* = 30 °C (black and grey, respectively; N = 101 and 136 rupture events after exclusion of the first bin; Poissonian counting error bars only shown at 1.1 pN for the sake of clarity). Data were fitted by single-exponential distributions to yield characteristic times, 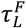, equal to 38.3 ± 5.4 and 5.16 ± 0.57 s (± SE). Inset, same plots for a *RBD*_*Wuhan*_/*ACE*2 complex (blue and cyan, respectively; N = 150 and 138) leading to 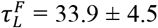 and 11.3 ± 1.3 s. **(F)** Force-dependence for the dissociation rate constant, 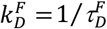, measured on *RBD*_*Wuhan*_ /*ACE*2 (blue), and 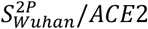 (black) at various temperatures. Data were fitted to the Bell model (Eq. 3 in Materials and Methods) to provide the dissociation rate constant at zero-force, *k*_*D*_, and the distance to the transition state, *x*^‡^ (table S3). **(G)** Temperature-dependence of *k*_*D*_ for *RBD*_*Wuhan*_ (blue) and 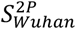 (black). Data were fitted to the Arrhenius equation (Eq. 4) to yield the activation energies, *E*_*D*_ (table S3).

The dissociation at slow shear rates or low forces is also important for viral infection (*16*). While HS-FS covers a wide range of loading rates, it has a limited force sensitivity (≳10 pN). Thus, to access lower forces, we employed MT (*51*) with the same ybbR constructs engrafted on modular junctured-DNA (J-DNA) scaffolds (Fig. 3D) (*52*). This procedure enabled the repeated probing of the dissociation of single *RBD*_*Wuhan*_ or 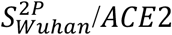 complexes at forces as low as ∼1 pN. The histogram of dwell time in the looped conformation (bound complex) at each applied force followed an exponential distribution with characteristic time 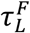 (Fig. 3E). By averaging 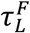 across different J-DNA scaffolds and correlating it to the dissociation rate constant under force, we characterized the energy landscape by fitting the Bell model (*53*) (Fig. 3F). We obtained in this manner *k*_*D*_ ∼(23 ± 2)×10^-3^ s^-1^ and *x*^‡^∼0.8 nm for *RBD*_*Wuhan*_ and 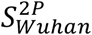 (Fig. 3F, table S3), in fair agreement with the values derived from HS-FS experiments. Additionally, the *k*_*D*_ values obtained at different temperatures and fitted with the Arrhenius equation led to activation energies *E*_*D*_, that were similar for *RBD*_*Wuhan*_ and 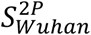 (Fig. 3G, table S3), again confirming that the dissociation pathway is not impacted by the integration of the *RBD* within *S* trimers, as suggested by previous SMFS data (see supplementary text S2) (*11*–*13, 15*–*17, 54*).

We then estimated the association kinetics by measuring the adhesion probability *P* at different contact times between an AFM tip functionalized with *ACE*2 and a surface coated with the *RBD* or the *S* trimer (Fig. 4A) (*55, 56*). *S* trimers appeared ∼2 to 4 times slower to bind to *ACE*2 (0.17-0.33 s) than isolated *RBD* (0.08 s). However, the interpretation of these results requires too many hypotheses on the behavior of the system at the molecular scale, which consequently makes the extraction of quantitative data difficult (supplementary text S3). Therefore, to accurately quantify binding, we monitored the looping and unlooping of J-DNA scaffolds at very low, constant force (0.1 pN) while increasing concentrations of *ACE*2 in solution (Fig. 4B-C). As expected, the dwell times in the looped conformation, Δ*t*_*L*_, obeyed monoexponential distributions with characteristic times 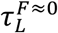 that did not depend on the *ACE*2 concentration and were similar for *RBD*_*Wuhan*_ and 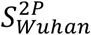 (Fig. 4D, F). It yielded *k*_*D*_ values (Fig. 4G, Table 2) in fair agreement with those derived above from MT-based SMFS experiments (table S3). In contrast, monoexponential fits to the histograms of the dwell times in the unlooped conformation, Δ*t*_*U*_, provided characteristic times 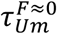 that increased with the *ACE*2 concentration – indicating that the formation of a complex between the two scaffolded components becomes more difficult as more *ACE*2 molecules from the solution occupy the *RBD*s (Fig. 4E-F). To be more quantitative, we studied the linear variation, as a function of *ACE*2 concentration, of the ratio between the time spent in the unlooped and in the looped conformation (*57*). In this way, we obtained the dissociation equilibrium constants for the complex of *ACE*2 with *RBD*_*Wuhan*_ and with 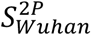 (Fig. 4H and Eqs. 5-6 in Materials and Methods): an *RBD* within the *S* trimer showed a ∼3.5-fold lower affinity than an isolated *RBD* (Fig. 4I, Table 2), which corroborates the BLI measurements (Fig. 2, Table 2). Considering the dissociation rate constant values, it translated into an association rate constant for an isolated *RBD* that is ∼2.5-fold higher than for an *RBD* within the *S* trimer.

**Fig. 4.**
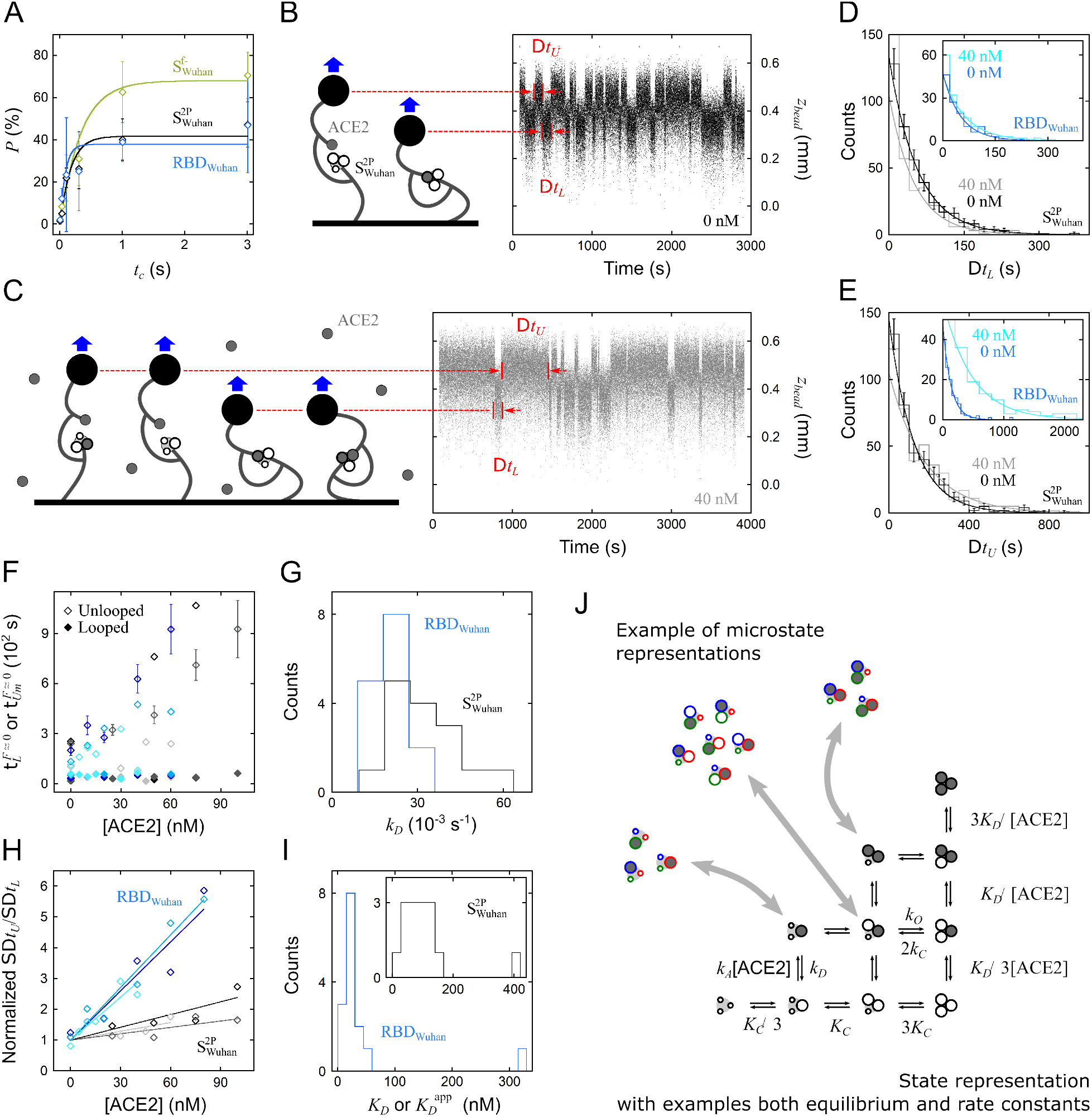
Single-molecule measurements of the kinetic and thermodynamic parameters for *ACE2* binding to the *RBD*, either isolated or embedded within the *S* trimer. **(A)** Adhesion probability, *P*, as a function of the contact time, *t*_*C*_, between the AFM cantilever functionalized with *ACE*2 and the surface coated with either *RBD*_*Wuhan*,_ 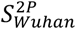 or 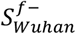 (blue, black, and kaki, respectively). Error bars correspond to SEM. Fits to the first-order kinetics provided the characteristic interaction time (*τ*_*U*_ = 0.08 ± 0.04 s for *RBD*_*Wuhan*_, 0.33 ± 0.13 s for 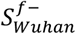, and 0.17 ± 0.06 s for 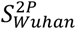). **(B, C)** Representative time traces obtained during the titration by soluble *ACE*2 of J-DNA engrafted with *ACE*2 and 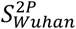, *ACE*2 being absent or present in solution at 40 nM. The applied constant force of 0.1 pN was low enough for the proteins on the scaffold to spontaneously dissociate and associate in *cis*; it was also high enough for the looped and unlooped conformations to be distinguished, and the corresponding dwell times, respectively Δ*t*_*L*_ and Δ*t*_*U*_, to be measured. **(D)** Examples of Δ*t*_*L*_ histograms collected on a 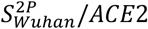 complex and fitted by single-exponential distributions to yield characteristic times for the looped conformation, 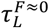, equal to 57.0 ± 4.4 and 50.9 ± 4.3 s (± SE) for [*ACE*2] = 0 and 40 nM (resp. black and grey, N = 262 and 187 rupture events). Inset, same plots obtained on a *RBD*_*Wuhan*_/*ACE*2 complex, with 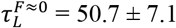 and 64.0 ± 8.0 s for [*ACE*2] = 0 and 40 nM (resp. blue and cyan, N = 83 and 109). **(E)** Examples of Δ*t*_*U*_ histograms collected on the same complex. For 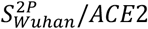, it gave characteristic times for the unlooped conformation 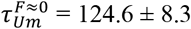 and 180.2 ± 12.6 s for [*ACE*2] = 0 and 40 nM (resp. black and grey, N = 256 and 300). For *RBD*_*Wuhan*_/*ACE*2 (inset) we obtain 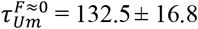 and 474.4 ± 66.0 s for [*ACE*2] = 0 and 40 nM (resp. blue and cyan, N = 90 and 98). **(F)** Variation, as a function of titrant concentration [*ACE*2], of 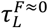 (full markers) and 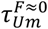 (open markers). Data are given for three representative scaffolds engrafted with *ACE*2 and *RBD*_*Wuhan*_ (blue to cyan) and three others with *ACE*2 and 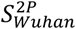 (black to grey). Each data point was obtained by monitoring a total of N= 58 to 310 events; error bars correspond to SE and, for clarity, they are only displayed for two examples. **(G)** Distributions of the dissociation rate constants obtained on individual scaffolds for *RBD*_*Wuhan*_/*ACE*2 and for 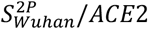 (resp. blue and black, 15 and 15 J-DNA, N = 73 to 764 dissociation events pooled per scaffold over the whole concentration range; Table 2). **(H)** Variation, as a function of the concentration in titrant, of the ratio between the total time spent in the unlooped conformation and the total time spent in the looped one. Linear fits with Eqs. 5-6 resulted in *K*_*D*_ = 17.5 ± 1.1, 18.8 ± 1.9, and 21.5 ± 2.7 nM for the three exemplary scaffolds engrafted with *ACE*2 and *RBD*_*Wuhan*_ (blue to cyan) and in 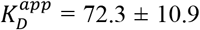, 104.8 ± 24.1, and 145.3 ± 28.7 nM for the three other ones with *ACE*2 and 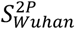 (black to grey). For each scaffold, both the data points and the fit have been renormalized by the fit value at [*ACE*2] = 0 nM, making it easier to visualize the dissociation equilibrium constant as the inverse of a slope (fig. S9 for the non-normalized data). **(I)** Distributions of the dissociation equilibrium constants obtained on individual scaffolds for *RBD*_*Wuhan*_/*ACE*2 (blue, 15 J-DNA) and for 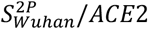 (inset, black, 15 J-DNA; Table 2). (**J**) Schematic description of all possible states for the *S* trimer and for its complexes with either one, two, or three *ACE*2 molecules, as well as of all possible transitions. The microstates are only represented for 3 states and similarly, the equilibrium and rate constants are only explicited for a subset of the reactions. *K*_*D*_ is the dissociation equilibrium constant of the *RBD*/*ACE*2 complex and *k*_*A*_ and *k*_*D*_ the corresponding association, and dissociation rate constants, respectively.

### Conformational selection model

All our experimental datasets, drawn from different experiments and methods, are coherent and converge towards a single model. Our results show that each *S* protomer (*S*^*protomer*^) acts independently and can either be open 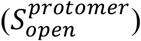 or closed 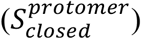, and associate to *ACE*2 according to the reaction network of Fig. 4J. To account for that, we propose a conformational selection model describing a serial, two-step mechanism. First, the *S* protomer in the trimer opens and, second, it binds to *ACE*2:

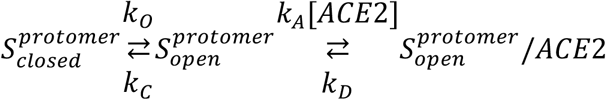

From the experimental point of view, it appears as a single-step process because the relaxation time characterizing the *RBD* conformational transition (*k*_*O*_ + *k*_*C*_) ^−1^ ∼0.7 s is much shorter than the time characterizing the binding-unbinding of *ACE*2 on the open *RBD* (*k*_*A*_[*ACE*2] + *k*_*D*_)^−1^ ≳ 13 s over the whole used concentration range (Tables 1 and 2). Therefore, the first step is always at equilibrium with respect to the second and this conformational selection scheme simplifies into:

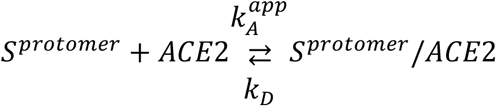

Since in the *S* trimer, each *RBD* has a probability *p*_*open*_ to be accessible to *ACE*2, it yields an apparent equilibrium dissociation constant 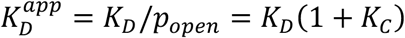 and, assuming unaffected dissociation rates, an apparent association rate 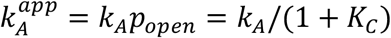 (supplementary text S4) (*58, 59*). Therefore, if the *RBD* opening probability is high, or the opening rate is much faster than the closing rate, 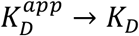, not affecting the binding. In contrast, if the opening probability tends to zero, or opening is much slower than closing, 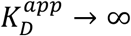, suppressing the binding. Importantly, because the binding is only modulated by *K*_*C*_, it does not depend on the *RBD* valency in the *S* trimer, making the model applicable to both trimers and protomers. The experimental *RBD* opening/closing rates presented in Table 1 resulted in *K*_*C*_∼2 for the 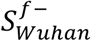 and 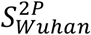 trimers, leading to 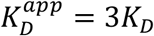, which quantitatively explains the values derived from 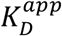 and *K*_*D*_ in both the BLI (ensemble) and MT (single-molecule) measurements presented in Table 2.

From these results, it appears that modulating the rapid opening and closing of the *RBD* tunes the affinity of the whole spike. This concept should be further evidenced with the 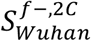 mutant in which a disulfide bond has been artificially inserted to impede opening. Indeed, according to the *K*_*C*_ obtained from HS-AFM imaging, this modulation should lead to an affinity to *ACE*2 up to 11 times lower than the one of 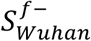 Previous structural and BLI works on similar *S* trimers mutant locked in the closed conformation are in agreement with our results (*24*).

Previous works showed that for the Omicron variant, the isolated *RBD* and the *S* trimer have a significantly higher binding affinity for *ACE*2 compared to the Wuhan variant, but some reports indicate all *RBD* in the closed state, which should result in a lower binding affinity (*23, 41, 60*). From our *K*_*C*_ values, we estimated a ∼6-fold reduction in *ACE*2 binding affinity for a *RBD* within the Omicron spike trimer compared to an isolated *RBD*. In theory, the higher probability for the Omicron spike to have all *RBD* in the closed state compared to the Wuhan spike should prevent infection. However, this is compensated by mutations in the *RBD* which enable it to complex *ACE*2 with higher affinity (*23, 41, 44, 61*). The advantage of this apparently neutral deal would be that an Omicron spike with lower probability of presenting *RBD* open is less susceptible to be neutralized by antibodies binding to the *RBD*. Therefore, our results partly explain the high immune evasion of the Omicron spike (*45, 62*).

Despite the enormous research effort undertaken since the emergence of COVID-19, the dynamics and functional mechanisms whereby SARS-CoV-2 enter their host have been challenging to explain. By combining molecular engineering with the ensemble and single-molecule techniques, we unveiled the multistep molecular mechanisms of the *S* trimer binding to *ACE*2. Our results show independence between conformations of the three protomers in the spike and a faster relaxation for the *RBD* opening and closing reaction than for *ACE*2 association and dissociation. We thus propose a conformational selection model in which the *RBD* open/close transition modulates the binding to *ACE*2, independently of the number of accessible binding sites and binding affinity. This model is especially significant for the Omicron variant, which presents an increased *RBD* closing equilibrium constant through mutations away from the receptor binding motif together with an increased *RBD*/*ACE*2 binding affinity constant (*41*). This would lead to a powerful mechanism to evade neutralizing antibodies that recognize the open *RBD*, while still binding to the host. Therefore, our results help explain the observed enhanced infectivity of Omicron and potentially of other SARS-CoV-2 variants of concern. Finally, the analysis of *RBD* opening and closing developed for HS-AFM imaging is adaptable to other techniques, such as cryo-EM or smFRET. We therefore provide a tool to predict the modulation of spike affinity by conformational changes and to test possible cooperative effects between protomers. Although not restricted to, we anticipate this approach to be useful for the study of other viral systems, *e*.*g*. future SARS-CoV-2 emerging variants and other human- or animal-borne coronaviruses (*63*). Furthermore, the experimental and conceptual framework we have developed should help in the conception of more effective vaccines and antibodies.

## Acknowledgements

We thank M. Backovic (UVS, Institut Pasteur, Paris) for logistics; M. A. Nash (Chemistry Dept., Univ. of Basel) for the gift of the sfp plasmid; P. England and the Biophysics Facility (Institut Pasteur) for access to the BLI; D. Joshi (RCAS, Academia Sinica, Taipei) for advises on PyMOL; P.-H. Puech and L. Limozin (LAI, Aix-Marseille Univ.) for insightful discussions; and J. Reguera (AFMB, Aix-Marseille Univ.) for critical reading of the manuscript.

## Author contributions

CG, TS, FAR, and FR supervised research. IF, AM, EB, and AS designed and produced the *ACE*2, *RBD*, and spike proteins. PS and FSu acquired the HS-AFM videos. PS and TB developed and applied the ImageJ macros to extract the conformational trajectories from HS-AFM videos. FR analyzed conformational trajectories. CG and FR developed the conformational selection model and implemented it for data analysis. IF, EB and PGC conducted BLI measurements. CV performed the AFM and HS-AFM SMFS experiments and analyzed data with FR. DK synthesized J-DNA and coupled them to the molecular partners. DK and JP realized MT measurements. FSt developed the software for analyzing the constant-force MT experiments. PS, CG, CV, and FR wrote the paper with the contributions of all the authors.

## Data and materials availability

All the materials and methods are detailed in the *Supplementary Materials*, including statistics. The code used for image processing is available on the GitHub repository (source code: https://github.com/centuri-engineering/ProtruDe/). Representative HS-AFM videos are provided in the *Supplementary Materials*. The raw data are available on request from the authors.

## Supplementary Materials

This PDF file includes: Materials and Methods, Supplementary Text S1 to S4, Figs. S1 to S9, Tables S1 to S3, Supplementary Videos (SV) information, References

Videos SV1 to SV20

