## Supplementary Information for "Modulation of SARS-CoV-2 spike binding to ACE2 through conformational selection"

**The PDF file includes:**

Materials and Methods
Supplementary Text
Figs. S1 to S9
Tables S1 to S3
References

**Other Supplementary Materials for this manuscript include the following:**

Videos SV1 to SV20

**Materials and Methods**

### 1. Biomolecule preparation

#### 1.1 Constructs design

Sequences for the $ACE2$, $RBD$, and spike ($S$) trimer constructs were codon optimized, synthesized, and cloned in the pcDNA3.1(+) vector by Genscript (Rijswijk, The Netherlands).

The ectodomain of the human $ACE2$ receptor sequence (residues 19-615, Uniprot Q9BYF1) was cloned downstream of a mouse IgK signal peptide and followed by a thrombin cleavage site and in-tandem Hisx8 and ybbR tags.

The $RBD$ sequences (residues 331-528, Uniprot P0DTC2) were cloned downstream of a mouse IgK signal peptide and followed by a thrombin cleavage site and in-tandem Hisx8, Strep, and ybbR tags. The Wuhan-type protein $RBD_{Wuhan}$ carried the V367F mutation (*1*), a substitution whose impact on our measurements is assumed to be minimal because it is located far from where the complex with $ACE2$ is formed.

Two wild-type spike expression plasmids were generated, comprising residues 1-1208 (Uniprot P0DTC2) followed by the foldon trimerization motif (YIPEAPRDGQAYVRKDGEWVLLSTFL) (*2*, *3*) and Hisx8, Strep, and ybbR tags. Wuhan-type $S$ proteins carried the V367F substitution. Two strategies were used to stabilize the metastable $S_{Wuhan}$ in the pre-fusion state. The first plasmid coded for $S_{Wuhan}^{2P}$, a spike stabilized by 2 proline mutations (K986P, V987P) (*3*, *4*) but in which the $^{682}$RRAR$^{685}$ furin cleavage site was unaltered. This construct has been widely used to comprehend antibody-mediated neutralization for the development of drugs and vaccines (*5*). The second plasmid coded for $S_{Wuhan}^{f-}$, a mutant stabilized by replacing the furin cleavage site with the $^{682}$GSAS$^{685}$ flexible linker (*3*). The latter plasmid was next used as a template to generate the vectors coding for the two cysteine-locked mutants: $S_{Wuhan}^{f-,2C}$ carrying mutations S383C and D985C, and $S_{WT}^{f-,2C,2P}$ carrying mutations S383C, D985C, K986P, and V987P. Independently, the synthetic gene coding for the Omicron spike (residues 1-1205, Genbank UFO69279.1) was designed with the furin site mutation described above as well as the C-terminal foldon motif followed by Hisx8, Strep, and Avi tags. This spike, denoted $S_{Omicron}^{f-,6P}$, was further stabilized with 6 proline mutations (F817P, A892P, A899P, A942P, K986P, V987P) relying on the previously reported HexaPro approach (*6*).

All $RBD$ and spike constructs used for BLI contained an Avi tag instead of a ybbR one.

#### 1.2 Protein expression and purification

The plasmids coding for recombinant proteins were transiently transfected in Expi293 cells (ThermoFisher Scientific, Waltham, MA) using the FectroPRO DNA transfection reagent (Polyplus, Illkirch, France) and following the manufacturer's instructions. After incubating the cells at 37 °C for 5 days, supernatants were recovered and concentrated using tangential flow filtration (TFF) cassettes (Vivaflow 200; Sartorius). Next, proteins were purified by affinity chromatography, $RBD$s and spikes on Strep-Tactin Superflow high-capacity cartridge columns (IBA, Göttingen, Germany), and $ACE2$ on a HisTrap Excel column (Cytiva, Velizy-Villacoublay, France). The eluted products were then concentrated and further purified by size-exclusion chromatography (SEC), $RBD$s and $ACE2$ on Superdex 200 Increase 10/300 columns (Cytiva) and spikes on Superose 6 Increase 10/300 columns (Cytiva), all having been pre-equilibrated with protein buffer, i.e. 10 mM Tris-HCl pH 8, 100 mM NaCl. SEC fractions were collected, pooled, concentrated, and stored at -80 °C until further use.

Note that for the AFM-based SMFS experiments, some intermediate buffer exchange was realized to condition proteins in phosphate buffer saline pH 7.4 (PBS).

### 2. High-speed atomic force microscopy (HS-AFM) imaging

#### 2.1 Sample preparation

A freshly cleaved muscovite mica disc with a diameter of 1.5 mm and a thickness of 0.1 mm (JBG-Metafix, Montdidier, France) was glued on a glass slide and used for the adsorption of proteins. The surface was first pre-incubated in a 10 mM $NiCl_2$ aqueous solution for 15 min, rinsed with water, and air dried. This approach resulted in a weakly positively charged surface, facilitating the partial immobilization of proteins while preserving their physiological functions and making it favorable for HS-AFM imaging. Then, we deposited a 2 µL droplet of *S* trimer dissolved at 1 – 2 µg/mL in imaging buffer, i.e. 10 mM Tris-HCl pH 7.4, 100 mM NaCl, 1 mM $MgCl_2$. After 30 min of incubation, the substrate was washed 10 times with imaging buffer to get rid of the unbound proteins.

#### 2.2 Data acquisition

Imaging was carried out in tapping mode on an SS-NEX HS-AFM equipped with ultrashort cantilevers having a resonance frequency of 600 kHz in liquid and a nominal spring constant of 0.15 N/m (USC-F1.2-k0.15, NanoWorld, Neuchâtel, Switzerland). During scanning, the free oscillation amplitude of the cantilever was set to 3 – 4 nm and the set point for feedback control was kept 20 % lower than this value. Images were acquired at a rate of 1 – 2 frames per second. Typically, the size of a pixel was 0.5×0.5 $nm^2$, while the size of a frame was 300×300 $nm^2$. At least 3 independent experiments were performed for each spike variant. All measurements were achieved at room temperature, i.e. 20 – 25 °C, in the imaging buffer.

#### 2.3 Data processing and analysis

An effort has been made in this work to enable semi-automatic quantitative analysis of HS-AFM videos. To this end, a pipeline has been developed to optimize image processing so that the many recorded images can be handled efficiently.

##### *2.3.1 Code availability*

All HS-AFM videos were semi-automatically processed and analyzed using in-house macros and plugins implemented in Fiji (ImageJ) (*7*). All the macros and plugins, together with the installation instructions, are available on GitHub repositories (source code: https://github.com/centuri-engineering/ProtruDe/). A step-by-step description of the process is provided below using a representative image extracted from an HS-AFM video, as depicted in figure S1.

##### *2.3.2 Pre-processing and cleaning*

The first routine (MacroPreProcessing) used the raw images of a video as input (fig. S1A) and, in one step, processed all the images of the video into a video with improved image quality (fig. S1B). More precisely, to start with, the noise was reduced by applying a median filter with a radius of 2 pixels. Then, the tilted background, which results from the angle made by the AFM tip with the surface during scanning, was corrected thanks to the ImageJ subtract background algorithm. This averages out all the pixels depending on the selected radius of a ball and subtracts this average value from the original image, thereby eliminating the spatial fluctuations of the background intensities. Here, we typically worked with a ball radius of around 50 pixels which is larger than the size of the $S$ protein.

A second routine (MacroThresholding) converted the pre-processed video (fig. S1B) into a binary video (fig. S1C) by setting a threshold value on the pixel intensity. An automated threshold employing either the Li or Triangle algorithm (*8*, *9*). A further process was used to discard objects smaller than the $S$ protein in the video. To do that, a threshold size was chosen based on the pixel size of the video and the size of the $S$ protein.

A third routine (MacroCleaningMask) used the binary, thresholded video (fig. S1C) and applied a dilation with a radius of 5 pixels, and further discarded all objects outside the main object. The resulting video was applied as a mask to the binary and pre-processed videos, resulting in a cleaned binary video and a cleaned video (fig. S1D).

*2.3.3 Spike alignment*

The alignment of $S$ protein involved the sequential use of the following ImageJ/Fiji plugins: AFM2_AlignBin, AFM2_FirstFrame, and AFM2_Reference. To accomplish this, the center of mass of the $S$ protein in the cleaned video (fig. S1D) was computed from the cleaned binary video generated by the MacroCleaningMask routine. This allowed the calculation of translation vectors that were subsequently applied to the cleaned video (fig. S1D), resulting in what we refer to as the 1st aligned video (fig. S1E). Next, image correlation was used to determine the optimal translation and rotation, enabling the alignment of all images with respect to a reference image. This step resulted in the generation of 2nd aligned video (fig. S1F). To further refine the process, the images within the resulting video were finally aligned with a reference derived by averaging all the images from the previous video (fig. S1G). This step was iterated until it provided a satisfactory outcome, which we denoted as the 3rd aligned video. Here, the stalk of the $S$ protein is indicated with a white arrowhead, whereas the bulbous head is marked with a yellow arrowhead (fig. S1H).

*2.3.4 Protrusion detection and characterization*

The MacroProtrusion routine transformed the 3rd aligned video (fig. S1H) into a binary video (fig. S1I). This aligned and thresholded video was then subjected to an erosion and dilation sequence using a disc with a radius of 5-10 pixels. The radius is chosen according to the pixel size of the video, taking into account the size of the protrusion ~5 nm. This morphological opening process yielded an opened video from which all protrusions having a size roughly smaller than the selected radius were removed (fig. S1J). Eventually, each opened video (fig. S1J) was subtracted from the corresponding binary one (fig. S1I) in order to keep only the protrusions (fig. S1K). Only the peripheral fragments of the $S$ protein that were larger than the expected radius of the protrusion remained present in the resulting binary video. The MacroDeleteROIProtrusion routine allowed the manual selection of regions of interest (ROI) on the bulbous head, to separate the protrusion (i.e. open $RBD$s) from the stalk and other artifactual bumps (fig. S1L). The MacroClassifyProtrusion routine extracted properties such as the number of detected $RBD$s per $S$ protein as well as the area and height of each $RBD$ from every image within a video, wherever required. These data were later used to generate conformational trajectories to analyze the transition kinetics of $RBD$ within $S$ protein.

The contour of the detected protrusion was visualized on the video using the MacroContourProtrusion routine. It used the aligned video (fig. S1H) and the binary video with ROI (fig. S1L) to produce a video with the contour of the ROI highlighted in cyan on each image. Finally, this video was combined side by side with the aligned video for visualization purposes (videos SV1-SV20).

*2.3.5 Transition kinetics analysis*

HS-AFM videos were used to generate conformational trajectories, also called transition time traces (Fig. 1B), that depict the opening and closing of the 3 $RBD$s using 4 different states for the $S$ trimer (0 to 3 $RBD$ open) over time. The mean relative occupancy of each conformational state and the associated standard error of the mean were computed from trajectories comprising a total of between 580 and 2780 images per $S$ trimer variant (table S2). An in-house developed Matlab code was used to compute the transition density plots (TDP), relying on between 230 and 930 transitions per variant. The dwell times distributions in each conformational state, relying on between 580 and 2780 images, were then used to globally fit monoexponential functions to the four most populated transitions and extract $k_C$ and $k_O$.

2.4. Simulated AFM images

To generate simulated AFM images, the BioAFMviewer software (*10*) was employed on cryo-EM structures of the $S$ trimers. The simulation parameters were set as follows: cone angle of 5° and tip radius of 1 nm. The following PDB files were utilized: 7DDD for 0 $RBD$ open (*11*), 7BNN for 1 $RBD$ open, and 7BNO for 2 $RBD$ open (*12*) (Fig. 1A). It provides estimates of the molecular configuration observed in the HS-AFM images.

**3. Bio-layer interferometry**

3.1 Sample preparation

$RBD$ and $S$ proteins were biotinylated using the BirA500:BirA biotin-protein ligase standard reaction kit (Avidity, Aurora, CO) according to the manufacturer's recommendations. After the buffer was exchanged for PBS, the biotinylated proteins were immobilized on streptavidin (SA) sensors (Sartorius, Aubagne, France) by immersion in solution at 50 nM ($RBD$) or 230 nM ($S$ trimer) for 600 s. The analyte solutions containing 200 nM to 1.56 nM $ACE2$ were prepared in PBS supplemented with 0.2 mg/mL BSA.

3.2 Data acquisition

Measurements were carried out on an Octet RED384 instrument (ForteBio, Fermont, CA) at room temperature. Association and dissociation were monitored for 150 s each. The sensor reference measurements were recorded from a sensor not loaded with either $RBD$ or $S$ trimer and dipped in the $ACE2$ solutions; the sample reference measurements were recorded from a sensor loaded with either $RBD$ or $S$ trimer and dipped in the assay buffer.

3.3 Data processing and analysis

Specific signals were calculated by double referencing, i.e. by subtracting the nonspecific signals obtained for both sensor and sample reference measurements from the signals recorded for the $RBD$ or $S$ trimer-loaded sensors dipped in the solutions of $ACE2$. Dissociation and association profiles were globally fitted on the instrument proprietary software assuming a 1:1 binding model. It provided first the dissociation rate constants, $k_D$, and next the association ones, $k_A$ or $k_A^{app}$. The dissociation equilibrium constants were finally obtained using either $K_D = k_D/k_A$ or $K_D^{app} = k_D/k_A^{app}$.

### 4. Atomic force microscopy-based single molecule force spectroscopy

#### 4.1 Sample preparation

All proteins ($ACE2$, $RBD$, and $S$ trimers) were engrafted via the ybbR tag added at their C-terminus allowing to control the pulling axes (*13*, *14*). They were engrafted on amine-modified AFM tips or glass coverslips through a 5 kDa PEG linker, which corresponds to a stretched length of around 32 nm. The tip/surface functionalization protocol was based on the previous publications (*14*, *15*) and the density of proteins was empirically adjusted to form single complexes. More precisely, cantilevers, either MLCT-Bio (Bruker, Billerica, MA), AC10 (Olympus, Tokyo, Japan), or AC40 (Olympus, Tokyo, Japan), were first rinsed in acetone and cleaned for 15 min in a UV Ozone reactor (ProCleaner; Bioforce Nanosciences, Virginia Beach, VA). The coverslips (1.5 or 6 mm, VWR, Radnor, PA) were rinsed in acetone, cleaned in a piranha solution ($H_2O_2/H_2SO_4$ 2:1) for at least 30 min, extensively rinsed with water, and dried. Next, all objects were silanized by immersing for 10 min in a 5 % ethanolic solution of 3-aminopropyldimethylethoxysilane (APDMES; ABCR, Karlsruhe, Germany). This was followed by rinsing in ethanol and water, drying with nitrogen, and baking for 30 min at 80 °C. The obtained amino-modified surfaces were subsequently incubated overnight at 4 °C in 0.1 M sodium borate pH 8.5-9, to yield $NH_3$ groups able to further react with carboxylates activated by N-hydroxysuccinimide. The following morning, the cantilevers and coverslips were functionalized by incubation for 1 h in a drop of 25 mM NHS-PEG-maleimide (5 kDa; Nanocs, New York, NYC) in sodium borate buffer, rinsed in water, passivated by incubation for 1 h in a drop of 250 mM NHS-PEG-methyl (MS(PEG)$_4$; ThermoFisher Scientific) in sodium borate buffer, and rinsed in water. Afterwards, the maleimide extremities of the PEG linkers were reacted 1 h with the thiol group of coenzyme A (CoA; Sigma Aldrich, Saint-Louis, MO) dissolved at 20 mM in 50 mM $NaHPO_4$ pH 7.2, 50 mM NaCl, 10 mM EDTA, followed by a new rinsing step in water. Then, 100 µL of each protein-ybbR construct at about 0.3 µM in PBS were mixed with 12 µL of in-house produced Sfp phosphopantetheinyl transferase (*13*) at 10 µM in 50 mM HEPES pH 7.4, 10 mM $MgCl_2$, and 12 µL of 100 mM $MgCl_2$. To conjugate the protein, 30 µL of this mixture was deposited onto surface. $ACE2$ was engrafted on the AFM tip and $RBD$ and $S$ trimers on the glass slides. We let the enzymatic reaction proceed for 1 h before samples were rinsed with phosphate buffer saline (PBS) pH 7.4 and stored in the same at 4 °C until further use.

#### 4.2 Data acquisition

SMFS measurements were achieved on a Nanowizard IV AFM (JPK BioAFM - Bruker Nano, Berlin, Germany) and an SS-NEX HS-AFM (Research Institute of Biomolecule Metrology, Tsukuba, Japan). Data were collected at room temperature, i.e. 20 – 25 °C, in PBS pH 7.4. Force-extension curves were collected by applying setpoints of 100–200 pN on 10×10 µm$^2$ lattices.

To obtain insight into the binding strength, the contact duration was set at 0.1s and the cantilever retraction velocity was varied, in random order, from 0.1 to 1000 µm/s. At least 300 force curves were acquired for each condition, on 3 different regions of each sample. Measurements were done at least in triplicate (8 independent pairs of cantilevers and coverslips for $RBD_{Wuhan}$, $S_{Wuhan}^{f-}$, and $S_{Wuhan}^{2P}$).

To obtain insight on the association kinetics, the AFM tip was approached and retracted at constant velocity, 2 μm/s, and the contact duration, $t_C$, was varied, in random order, from a set of predefined values varying from 0 to 3 s (*16*). At least 300 adhesion attempts were made for each condition. From these adhesion attempts, we calculated the fraction of binding events.

4.3 Data processing and analysis

Data processing and analysis were carried out using the JPK Data Processing software (JPK BioAFM - Bruker Nano) and an in-house developed tool written in Matlab (Mathworks Inc.). To generate the dynamic force spectra shown in Fig. 3C, rupture forces, $F_r$, were pooled by loading rate $\dot{F}$, and the obtained distributions were fitted with bimodal Gaussian functions to determine the most probable rupture force, $|F_r|$, and the associated standard error of the mean (SEM, figs. S6-S8). These values were next plotted as a function of the average $\dot{F}$ and were fitted using the expression derived by Cossio, Hummer, and Szabo (CHS), so as to determine the position of the transition state in space and energy, $x^‡$ and $\Delta G^‡$, and the dissociation rate constant, $k_D$ (*17*):

$$|F| = \frac{\Delta G^‡}{\mu x^‡}\left[1 - \left(\frac{k_B T}{\Delta G^‡}\ln\Lambda\right)^{\mu}\left(1 + \frac{\mu(3\mu - 2)\ln(\ln\Lambda)}{\ln\Lambda}\right)\right] \qquad \text{(Eq. 1)}$$

where,

$$\Lambda = k_D\left(\frac{\Delta G^‡}{k_B T}\right)^{2-3\mu} \exp\left(\frac{\Delta G^‡}{k_B T}\right)\frac{k_B T}{\hat{F} x^‡} e^{\gamma} \qquad \text{(Eq. 2)}$$

with $\gamma \approx 0.5772$ the Euler-Mascheroni constant and $\mu$ the kinetic ductility of the complex, the later parameter being fixed to 0.01.

To generate the curves of the binding probability over contact time, $P(t_C)$, shown in Fig. 4A, the average binding probabilities $P$, and the standard error of the mean, were computed at each contact duration $t_c$ from at least three independent experiments. These data were fitted to a first-order kinetic model $P = A(1 - \exp[-(t_C - t_0)/\tau_U])$, with $A$ a constant depending on the molecular density on the surface, $t_0$ the 10 ms instrument lag time, and $\tau_U$ a time constant characterizing the unbound cantilever status (*18*).

### 5. Magnetic tweezers-based nanomanipulation

5.1 Sample preparation

The symmetrical J-DNA forceps we relied on have been recently described (*19*) and were prepared using a slightly modified version of our previous protocol (*20*). They consisted of two linear dsDNA segments, the branches, connected by a third one, the leash. Their respective sizes were 1507, 1470, and 689 bp. The junctures with the leash divide the branches into tips and shanks. The tips were 48 bp-long and terminated with Nb.BbvCI-generated overhangs that are used for engraftment of the protein-oligonucleotide conjugates. The shanks had their extremities ligated to sticky ends, i.e. ≈ 1000 bp-long dsDNA fragments that are multiply-labeled with either biotin or digoxigenin, and were employed for respective attachment to the streptavidin-coated beads or to the anti-digoxigenin-coated surface.

The proteins carrying an ybbR tag at their C-terminus were conjugated to oligonucleotides labeled with CoA at their 5' end (Biomers, Ulm, Germany). Two different sequences were used, each complementary to a specific nine-base overhang present at one of the J-DNA tips: P1 = 5'-TTGTAAGAGC-3', P2 = 5'-TATATGAGGC-3'. To prepare the $ACE2$-P1 conjugate, 50 µg of CoA-P1 (11.9 nmol) were resuspended in 10 µL of water and combined with 60 µL of $ACE2$-ybbR at 127 µM in protein buffer (7.6 nmol). To prepare the $RBD_{Wuhan}$-P2 conjugate, 50 µg of CoA-P2 (11.9 nmol) were resuspended in 10 µL of water and combined with 60 µL of $RBD_{Wuhan}$-ybbR at 124 µM in protein buffer (7.4 nmol). To prepare the $S_{Wuhan}^{2P}$-P2 conjugate, 4.2 µg of CoA-P2 (1 nmol) were resuspended in 1 µL of water and combined with 45 µL of $S_{Wuhan}^{2P}$-ybbR at 13.5 µM in protein buffer (0.6 nmol in protomer). Each mixture was next supplemented with 1 µM Sfp (same preparation as above) plus 20 mM $MgCl_2$. After incubation for at least 2 h at 25 °C, the protein-oligonucleotide conjugates were purified by gel filtration chromatography using a Superdex 200 Increase column (GE Healthcare, Velizy Villacoublay, France) equilibrated in protein buffer. The concentration in products was estimated using the following theoretical values for the molar absorption coefficient at 280 nm: 151 720 $M^{-1}cm^{-1}$ for $ACE2$, 38 850 $M^{-1}cm^{-1}$ for $RBD_{Wuhan}$, and 134 030 $M^{-1}cm^{-1}$ for each $S_{Wuhan}^{2P}$ protomer.

The protein-oligonucleotide conjugates were ligated onto the J-DNA forceps using T4 DNA ligase (New England Biolabs, Ipswich, MA) in protein buffer supplemented with 1 mM ATP and 1 mM DTT. Reaction volumes of 50 µL containing each conjugate at 200 nM, J-DNA at 50 pM, and 200 U of T4 DNA ligase were incubated for at least 4 h at 16 °C. The assembled scaffolds were then aliquoted, frozen in liquid nitrogen, and stored at -80 °C.

The force-cycling experiments were carried out using glass flow cells that were either passivated with polystyrene and functionalized with anti-digoxigenin (*21*), or with PEG-linkers and functionalized with streptavidin (*22*). In the first strategy, J-DNAs were then attached to 1 µm-diameter magnetic beads coated with streptavidin (MyOne Streptavidin C1, ThermoFisher Scientific, Illkirch, France). In the second strategy, J-DNAs were attached to 1 µm-diameter magnetic beads (MyOne Tosylactivated, ThermoFisher Scientific) that have been functionalized in-house with anti-digoxigenin (*22*). The constant-force experiments were only employed in this last approach.

5.2 Data acquisition

All measurements were performed in 10 mM Na-HEPES pH 7.4, 150 mM NaCl, 0.1 % Tween 20, and 0.5 mg/mL BSA. Experiments were carried out on home-built magnetic tweezers setup running under the PicoJai software suite (PicoTwist, Fleurieux-sur-l'Arbresle, France) (*19, 23–25*). Beads were tracked with ≈ 5 nm accuracy in the three spatial dimensions and at 31 Hz. The end-to-end extension of each J-DNA scaffold was calculated from the position of the magnetic bead above the surface, using a bead stuck on the slide as a fiduciary marker of surface drift. The force was determined from the position of the magnets above the sample, relying on a calibration curve established on 17 kbp-long supercoilable dsDNA molecules – see (*25*) for an example of the procedure.

To obtain an insight on the dissociation kinetics and to map the energy landscape, force-cycling experiments were performed either at 25, 30, or 37 °C. The force applied on the magnetic bead was repeatedly alternated between a low and a high value, respectively 0.001 and either $F$ = 1.1, 3.9, 6.8, or 9.6 pN (Fig. 3D). The time spent at nearly 0 pN was equal to 13 s and the time spent applying force $F$ to the complexes ranged between 55 and 155 s, being shorter when pulling stronger.

To measure the dissociation equilibrium constants, titration experiments were performed at 30 °C, with the force kept constant at around 100 fN, a value for which the energy barrier towards association is surmountable. Free $ACE2$ receptor was added at concentrations varying from 0 to 160 nM; hence, it competed with the $ACE2$ molecule engrafted on the scaffold and prevented to a certain extent its association with the viral counterpart located at the other tip (Fig. 4C). For each viral partner, at least three independent titration experiments were realized, several beads being monitored in parallel during each of them.

5.3 Data processing and analysis

Results from the force-cycling experiments (Fig. 3D) were analyzed using custom routines in the Xvin software suite (PicoTwist), as described in our previous paper (*20*). More precisely, 25 to 230 rupture events were observed for each scaffold and for each condition. In the $RBD_{Wuhan}$ case, 5 to 15 scaffolds were monitored per condition, that is 645 to 1822 events in total. As far as $S^{2P}_{Wuhan}$ was concerned, we had 2 to 9 scaffolds per condition and 226 to 1171 events in total. As a rule, we excluded from analysis every event for which unbinding did not occur within the same cycle as the one in which binding was first observed. Dwell times in the looped conformation, $\Delta t_L$, were represented in histogram form with error bars equal to the square root of the number of events in each bin, which corresponds to the Poissonian counting error. After removal of the first bin (for which missed events may lead to undercounting), single-exponential, weighted, least squares fits were performed so as to yield, for each scaffold, a characteristic time $\tau^F_L$ (± SE) (Fig. 3E). Next, for each force, we first averaged the results obtained on the different J-DNAs and determined the standard error - as it was done everywhere else in our analysis, we here corrected the variance to remove the bias introduced by the small number of beads (*26*). Second, we computed the logarithm of the dissociation rate constant under force $F$ as $-\ln\langle\tau^F_L\rangle$ and propagated the errors. Finally, a weighted, least squares fit to the linearized Bell equation:

$$\ln k^F_L \ = \ln k_D + x^{\ddagger}/k_B T \qquad \text{(Eq.3)}$$

provided the dissociation rate constant at zero-force, $k_D$, and the distance to the transition state, $x^{\ddagger}$ (± SE) (Fig. 3F). Activation energies, $E_D$, were calculated using the Arrhenius equation:

$$\ln k_D \ = \ \ln A - E_D/RT \qquad \text{(Eq.4)}$$

and two $\ln k_D$ values obtained at two different temperatures; errors being propagated accordingly.

Time traces from constant-force titrations (Fig. 4B-C) were processed relying on an in-house program that enables to identify both looped and unlooped conformations thanks to hidden Markov modeling (*25*). Briefly, data were first cleaned from artefactual detection points, corrected for long-term drift to obtain a baseline as flat as possible, and submitted to a 1.6 s sliding average to reduce dispersion. Then, we used the hmmlearn library based on the sckit-learn Numpy Python package. The routine, which belongs to the Viterbi group of hidden Markov model learning algorithms, was trained on a limited portion of the time trace (typically 1000 s) to characterize the two conformations (i.e. the two HMM states). Once the learning was completed, a conformation (i.e. either looped or unlooped) was assigned automatically to every time point of the whole acquisition, which typically lasted 3 to 28 h. From the resulting binary signal, it was finally an easy task to identify the transitions and to compute the looped and unlooped dwell times, $\Delta t_L$ and $\Delta t_U$, accordingly.

In all, 51 to 964 dissociation (or equivalently association) events were observed for each scaffold and each condition, 17 complexes being monitored in the $RBD_{Wuhan}$ case and 22 in the

$S_{Wuhan}^{2P}$ one. Dwell times histograms displaying less than 50 events after removal of the first bin were not considered. Similarly, for the $\sum \Delta t_U / \sum \Delta t_L$ computations, we discarded time traces for which less than 50 looping events were observed. In addition, scaffolds that endured less than 3 $ACE2$ concentration conditions or presented a non-physical outcome (such as a negative dissociation equilibrium constant) were excluded: it corresponded to 2 J-DNA scaffolds for $RBD_{Wuhan}$ and 7 for $S_{Wuhan}^{2P}$.

Dwell times in the looped and unlooped conformations were represented in histogram form and fitted by monoexponential functions as described above for the analysis of the force-cycling measurements (Fig. 4D-E). It yielded the characteristic times $\tau_L^{F\approx 0}$ and $\tau_{Um}^{F\approx 0}$ (Fig. 4F). The second parameter is in fact only an approximation of the slowest decay of the true $\Delta t_U$ distributions, which is biexponential in the $RBD_{Wuhan}$ case (*25*) and certainly with many more components in the $S_{Wuhan}^{2P}$ case. Since, for each J-DNA, $\tau_L^{F\approx 0}$ is expected not to vary with $[ACE2]$ we pooled the $\Delta t_L$ dwell times collected at all concentrations and by exponential fitting of the resulting histogram obtained a $\tau_L^{F\approx 0}$ value per scaffold (Fig. 4F). Inversion next yielded a $k_D$ value per scaffold. Finally, the data obtained on all J-DNA were represented in histogram forms (Fig. 4G) and used to compute a mean ± standard error for each complex population (Table 2). Please note that in the $S_{Wuhan}^{2P}$ case we excluded one outlier with a very large $k_D$.

For the $RBD_{Wuhan}/ACE2$ complexes, ratios between the total time each scaffold spends in the unlooped conformation, $\sum \Delta t_U$ and the total time it spends in the looped one, $\sum \Delta t_L$, were analyzed as a function of the concentration in titrant (*25*):

$$\sum \Delta t_U / \sum \Delta t_L = K_U + K_U/K_D \ [ACE2] \quad \text{(Eq.5)}$$

with $K_D$ the complex dissociation equilibrium constant, $K_U = K_D/C'_{eff}$ the scaffold unlooping equilibrium constant, and $C'_{eff}$ the effective concentration at the DNA scaffold tips. Thus, linear fitting provided a slope $a$ and an intercept $b$, parameters which were further used to compute a $K_D$ value for each complex (Fig. 4H and fig. S9):

$$K_D = b/a \quad \text{(Eq.6)}$$

These results were then plotted as a histogram (Fig. 4G) and the distribution mean ± standard error was computed after exclusion of the outlier with a very large $K_D$ (Table 2). The same procedure was followed for the J-DNA engrafted with $ACE2$ and $S_{Wuhan}^{2P}$, except that, in the above formulas, $K_D$ was replaced by $K_D^{app}$ to account for $RBD$ opening and closing and $K_U$ was equal to $K_D^{app}/3C'_{eff}$, the factor 3 being related to the spike trimeric nature. Finally, association rate constants were derived as $k_A = k_D/K_D$ and $k_A^{app} = k_D/K_D^{app}$ and errors propagated (Table 2).

### Supplementary Text

S1. Conformational states of $RBD$ in $S$ trimers

*S1.1 Advantages of HS-AFM in comparison with cryo-EM and smFRET*

While cryo-EM offers high-resolution structural insights into the possible conformational states of the $S$ trimer, this provides static pictures of $RBD$ in the open and closed states. Hence, the information about real-time conformational changes is lacking. In addition, a recent study has highlighted the significant sensitivity of buffer composition in discerning the conformational states of the $S$ trimer, suggesting the need for complementary biophysical approaches to validate the structural models (*27*).

Some studies explored the real-time dynamics of the $RBD$ conformational states in solution. In smFRET experiments, two sites (on the $RBD$ and SD1 regions, positions 427 and 556, respectively) were labeled to track the relative movement of $RBD$ with millisecond time resolution (*28*, *29*). However, it is important to note that this method is constrained to the sites where fluorophores are attached and hence the complete picture of $RBD$ is missing. Moreover, in smFRET experiments, it is difficult to discern between the simultaneous opening of the three $RBD$s within the $S$ trimer since these smFRET measurements do not provide a view of the full protein structure and require structural data to guess what the different dynamic states are. Depending on the site chosen for fluorophore engraftment, the FRET signal may be sensitive to the conformation of the individual labeled protomer (*29*) or to the conformation of the whole spike (*28*). The last case is thus potentially more informative; however, connecting each energy transfer level to a given structure is complex, which may obscure the interpretation of the collected time traces. On the contrary, in the latter case linking FRET states to $RBD$ positions is quite straightforward but it may be difficult to infer from the data any global mechanism at the spike level. Therefore, HS-AFM direct visualization of all 3 $RBD$s stands as the only technique that can detect cooperative protomer motion.

Recently, HS-AFM on unlabeled $S$ trimers unveiled the dynamic conformations of the stalk and of the $RBD$s, resolving up to two $RBD$s in the open conformation and revealing the potential of this technique (*30–32*). However, it is worth noting that the kinetic rate constants governing $RBD$s opening and closing were not reported.

*S1.2 Comparison of our HS-AFM results with cryo-EM and smFRET data*

It can be informative to compare our $RBD$ state occupancy findings with published classification data obtained in cryo-EM (table S1). In many cases, for the Wuhan variant, only 0-$RBD$ open and 1-$RBD$ open $RBD$ states were detected, with $S_0$ populations ranging from 0 to 94 %. However, the majority of studies reported 0 $RBD$ open state populations between 40-60 %, in good agreement with our results. Applying the rough estimate approach used for our HS-AFM, we calculated that the closing equilibrium constant ($K_c \equiv \Sigma RBD_{closed}/\Sigma RBD_{open} = \sum_i (3-i) \times S_i / \sum_i i \times S_i$) determined from the cryo-EM particle classification varied between 1 and 49, most of them clustering between 3 and 5, consistent with our results. These $K_C$ values are similar but slightly larger than ours (around 2.5-3.5). However, such structural studies hardly report in a quantitative manner on what is present in the sample in solution and at equilibrium (*33*). This is evident from the significant variations observed across different articles and from the possible artifacts mentioned in (*27*). To notice, using HS-AFM, we evidenced the 2 $RBD$ open state and, although rare, the 3 $RBD$ open state. Both states were rarely detected by cryo-EM on

Wuhan variants although 2 $RBD$ open states have been reported, and both 2 $RBD$ and 3 $RBD$ open states have been observed in the presence of $ACE2$, and on other variants (table S1). Of interest, the comparison between 2P and non-2P Wuhan data showed no significant difference in most of the works, in agreement with our results. In the case of the Omicron variant, some cryo-EM works reported only $RBD$ open states, while others reported only closed $RBD$, in better agreement with our results.

The rate constants we found for $RBD$ conformational changes can also be compared with the data reported from smFRET measurements (*28*, *29*). smFRET found slightly faster kinetics, with opening ~4 to 5 times faster and closing ~1.5 to 4 times faster. More importantly, smFRET works have reported up to four different FRET states, which are difficult to assign to specific $RBD$ conformations. This could be interpreted as intermediate states with faster kinetic transitions that the one leading to full opening, intermediate states that we could not evidence in our work because of possible temporal and spatial resolution issues (*28*, *34*). Apart from additional data processing and state assignment biases, other explanations for this difference could include interactions with surfaces, labeling effects, or simply non-identical experimental conditions (e.g. temperature). Importantly, the qualitative fact that single opening and closing characteristic times were observed in spike including individually labeled protomer supports our observation of independent conformational changes (*29*).

S2. Dissociation Kinetics

Our results are in reasonable agreement with previous data from both single molecule and ensemble measurements and discrepancies can easily be explained by variation in the experimental conditions (*3*, *32*, *35–43*).

Some earlier AFM-based SMFS measurements (*35*, *37*) at low loading rates (<$10^5$ pN/s) reported higher rupture forces (40-70 pN) than those we observed (20-50 pN). This could be due to various reasons, for instance, different data analysis approaches and more likely, different pulling geometries. Indeed, the effect of the pulling direction on rupture forces has been reported for various receptor-ligand or protein-protein systems (*20*, *44*, *45*). While we used site-directed engraftment, through the C-terminal ybbR tag, these previous studies relied on molecules immobilized with random orientation. In line with this remark, other AFM-based SMFS measurements on the $RBD/ACE2$ attached by the C-terminus (thanks to enzymatic ligation or through NTA/his-tag) reported force values in better agreement with our results (20-60 pN) (*32*, *38*). Finally, a work using the biomembrane force probe revealed a catch-bond response at forces below 10 pN that we did not observe (*39*); however, the experimental systems are quite different.

The energy landscape parameters obtained in previous studies are in relatively good agreement with our results, despite the use of different models. AFM-based SMFS works reported distances to the transition state of ~0.4-2.0 nm for Wuhan type $RBD$ and $S1$ subunit binding to $ACE2$, within the range of our results (*32*, *35*, *37*, *40*, *42*, *46*). Interestingly, Bauer and coworkers (*42*) found much larger values for the transition state distance using MT and site-directed pulling. However, in their case, the $ACE2$ molecule was pulled from the N-terminus, while we pulled from the C-terminus. Again, pulling direction may explain the important change in the results, as reported for other receptor/ligand complexes (*44*).

The dissociation rate constant $k_D$ obtained from our HS-FS and MT experiments for Wuhan type $RBD$ and $S$ trimer (~$10^{-2}$ s$^{-1}$) is in good agreement with previous studies by AFM (*35*, *40*), MT (*42*), BLI (*47*, *48*), and SPR (*45*, *49*, *50*), while other works reported slightly higher values

(~5 to 8×10$^{-2}$ s$^{-1}$) using AFM (*37*, *38*, *42*, *46*) or SPR (*51*). As explained above, these differences can reflect differences in the protein orientation (being nonspecific (*37*, *46*) or opposite to the one we used (*42*) or can be induced by the tags used (use of biotinylated $ACE2$ (*51*) or mCherry-$ACE2$ (*38*)). Another possible explanation for these differences is temperature, which importantly affects $k_D$ (*48*, *50*).

S3. Association Kinetics

According to our AFM data, the characteristic time ($\tau_U$) for $ACE2$ to bind its viral partner is equal to 0.08 ± 0.04 s for $RBD_{Wuhan}$, 0.33 ± 0.13 s for $S^{f-}_{Wuhan}$, and 0.17 ± 0.06 s for $S^{2P}_{Wuhan}$. For monovalent interactions, the association rate constant can in principle be computed knowing $C_{eff}$, the effective concentration of the molecules facing each other: $k_A = 1/\tau_U\, C_{eff}$ (*16*). For $S$ trimers, the relation would transform into $k_A^{app} = 1/3\tau_U\, C_{eff}$ with the "app" superscript (for apparent) indicating that we are referring to a $RBD$ belonging to an $S$ protomer in which conformational changes take place, and with the factor 3 accounting for the trimeric nature of the spike. Estimating $C_{eff}$ can be done by considering that the partners, which are attached to surfaces through linkers, explore a hemisphere (*16*, *18*). More sophisticated 2D models can also be considered that assume the immobilized state of the receptor and ligand molecules (*52*). Yet, both strategies required either strong hypotheses on the protein grafting geometry or unknown protein densities, so we decided not to report on any association rate constants based on our AFM measurements.

S4. Computation of the relation between the equilibrium constants

The dissociation equilibrium constant for the reaction

$$S^{protomer}/ACE2 \rightleftarrows S^{protomer} + ACE2$$

is by definition given by

$$K_D^{app} = \frac{[S^{protomer}][ACE2]}{[S^{protomer}/ACE2]}$$

In this scheme, $S^{protomer}$ account for both the open and closed conformations, $S^{protomer}_{open}$ and $S^{protomer}_{closed}$ respectively. Furthermore, $S^{protomer}/ACE2$ is equivalent to $S^{protomer}_{open}/ACE2$ since binding can only occur when the $RBD$ is open.

Using the conservation of matter, one has

$$[S^{protomer}] = [S^{protomer}_{open}] + [S^{protomer}_{closed}]$$

which simplifies in

$$[S^{protomer}] = (1 + K_C)[S^{protomer}_{open}]$$

when considering the closing equilibrium

$$S^{protomer}_{open} \rightleftarrows S^{protomer}_{closed}$$

and its equilibrium constant $K_C$.

On the other hand, we have $[S^{protomer}/ACE2] = [S^{protomer}_{open}/ACE2]$. It then yields by substitution to

$$K_D^{app} = (1 + K_C)\frac{[S_{open}^{protomer}][ACE2]}{[S_{open}^{protomer}/ACE2]}$$

in which the fraction can be readily identified to $K_D$, the dissociation equilibrium constant of the binding of $ACE2$ to an available $RBD$

$$S_{open}^{protomer}/ACE2 \rightleftarrows S_{open}^{protomer} + ACE2$$

The sought-after result is consequently obtained as

$$K_D^{app} = (1 + K_C)K_D$$

### Supplementary figures

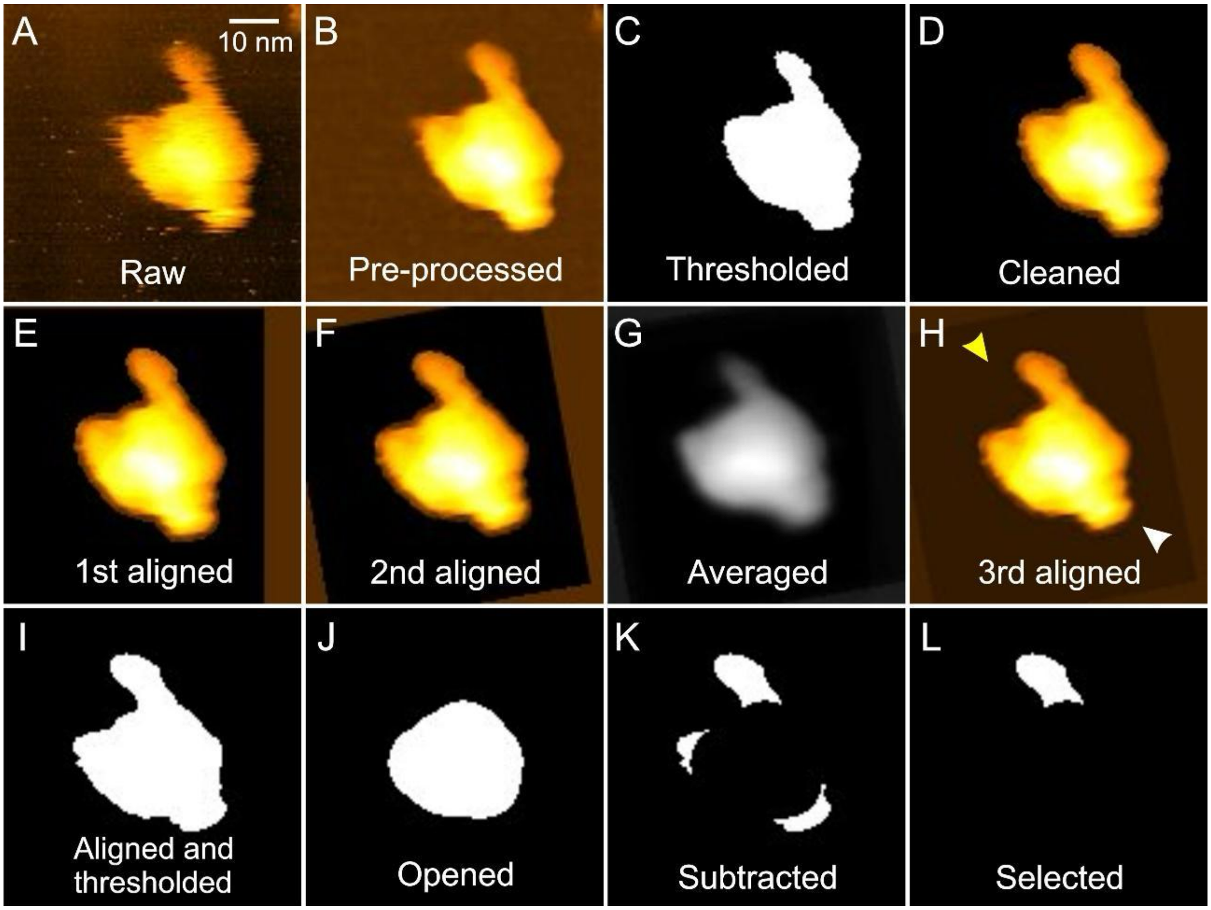


**Fig. S1.** Example of sequential HS-AFM image processing using ImageJ-based macros that lead to the detection of the protrusions on $S_{Wuhan}^{2C,2P}$. The white and yellow arrows on panel H indicate the stalk and the bulbous head of the spike, respectively. The x-y scale is shown in the top-right corner of the first image and is the same for other images.

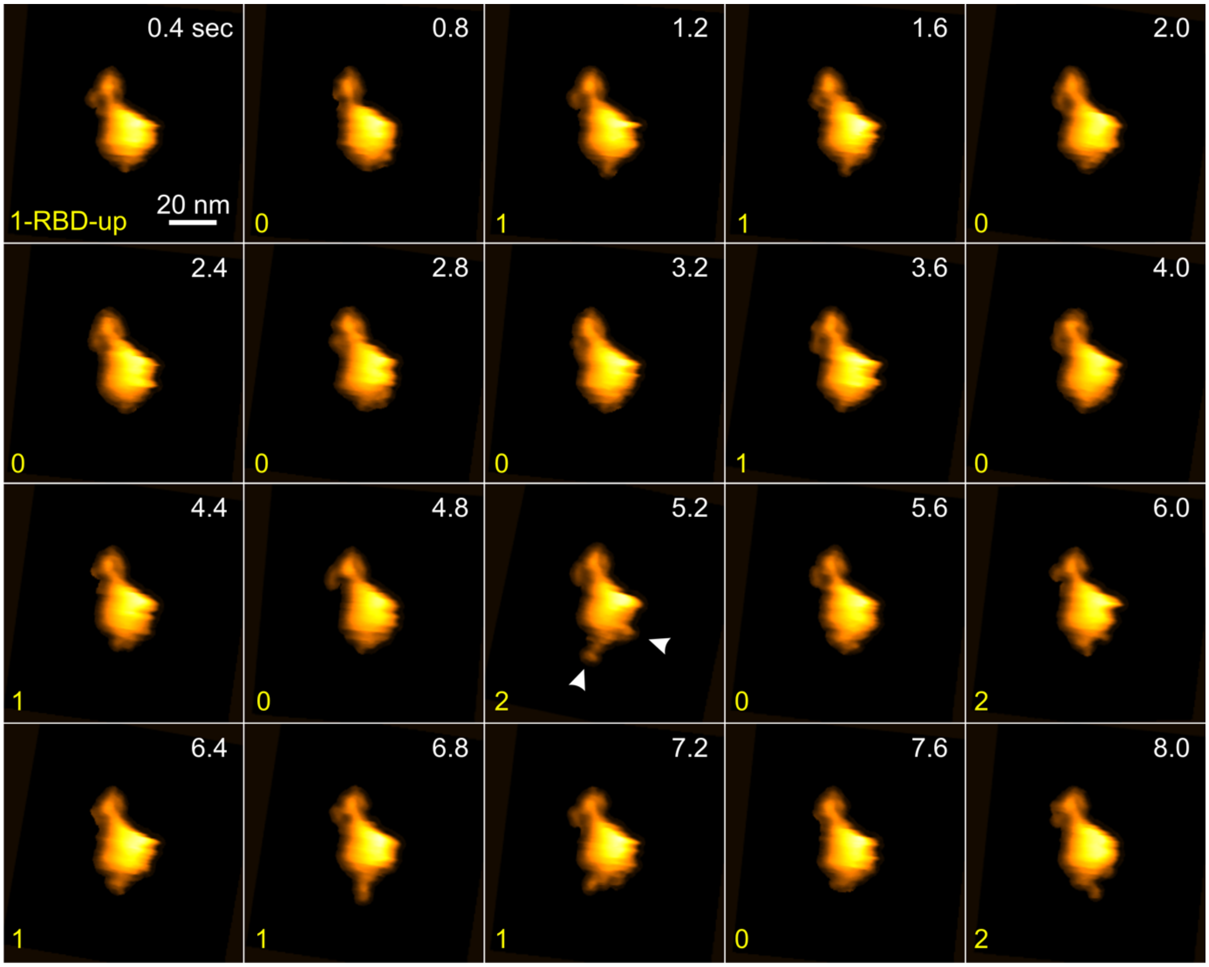


**Fig. S2.** Real-time $RBD$ conformations in $S^{f-}_{Wuhan}$. Consecutive images from an HS-AFM video depicting $S^{f-}_{Wuhan}$ in various states (as indicated in the bottom left corner of the images). The opening of $RBD$s is indicated with white arrowheads in one of the images at 5.2 s. The acquisition rate of the video was 2.5 fps. These images were processed with a set of user-written macros (see Materials and Methods) to reduce noise and remove small particles present in the background.

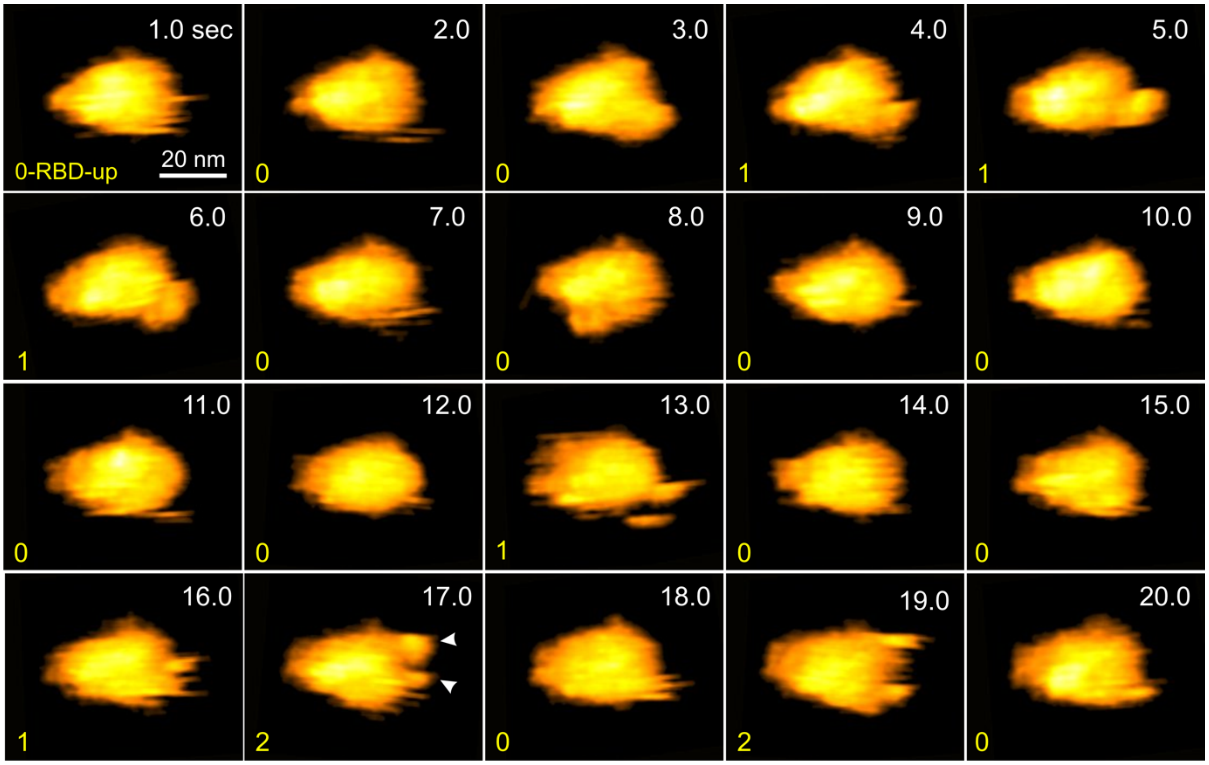


**Fig. S3.** Real-time $RBD$ conformations in $S^{2P}_{Wuhan}$. Consecutive images from an HS-AFM video depicting $S^{2P}_{Wuhan}$in various states (as indicated in the bottom left corner of the images). The opening of $RBD$s is indicated with white arrowheads in one of the images at 17.0 s. The acquisition rate of the video was 1.0 fps. These images were processed with a set of user-written macros (see Materials and Methods) to reduce noise and remove small particles present in the background.

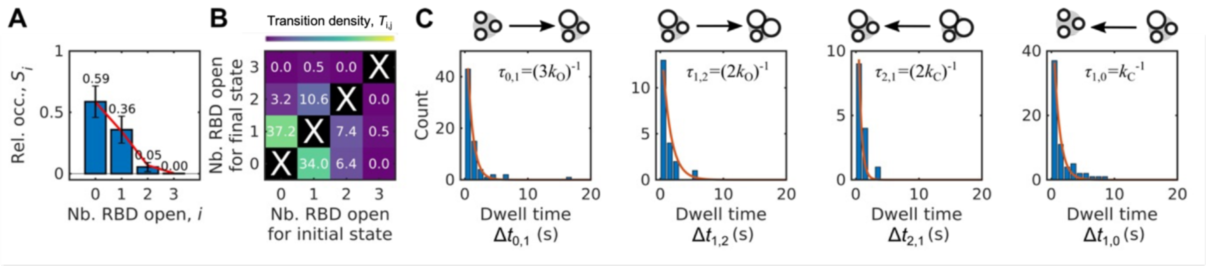


**Fig. S4.** Conformational dynamics of $RBD$ opening and closing in $S_{WT}^{f-,2C,2P}$ (A) Relative occupancy histogram (mean±SEM, blue bars) and associated fits to the binomial distribution (red lines) for the four $S_i$ states, $i$ indicating the number of opened $RBD$s. (B) Transition density plot (TDP) showing the fraction of transition events between the different states. (C) Distributions of the $\tau_{i,j}$ dwell times for the four most populated $S_i$ to $S_j$ transitions. Red lines show global fits of exponential decays (see Table 1 for results and table S2 for statistics).

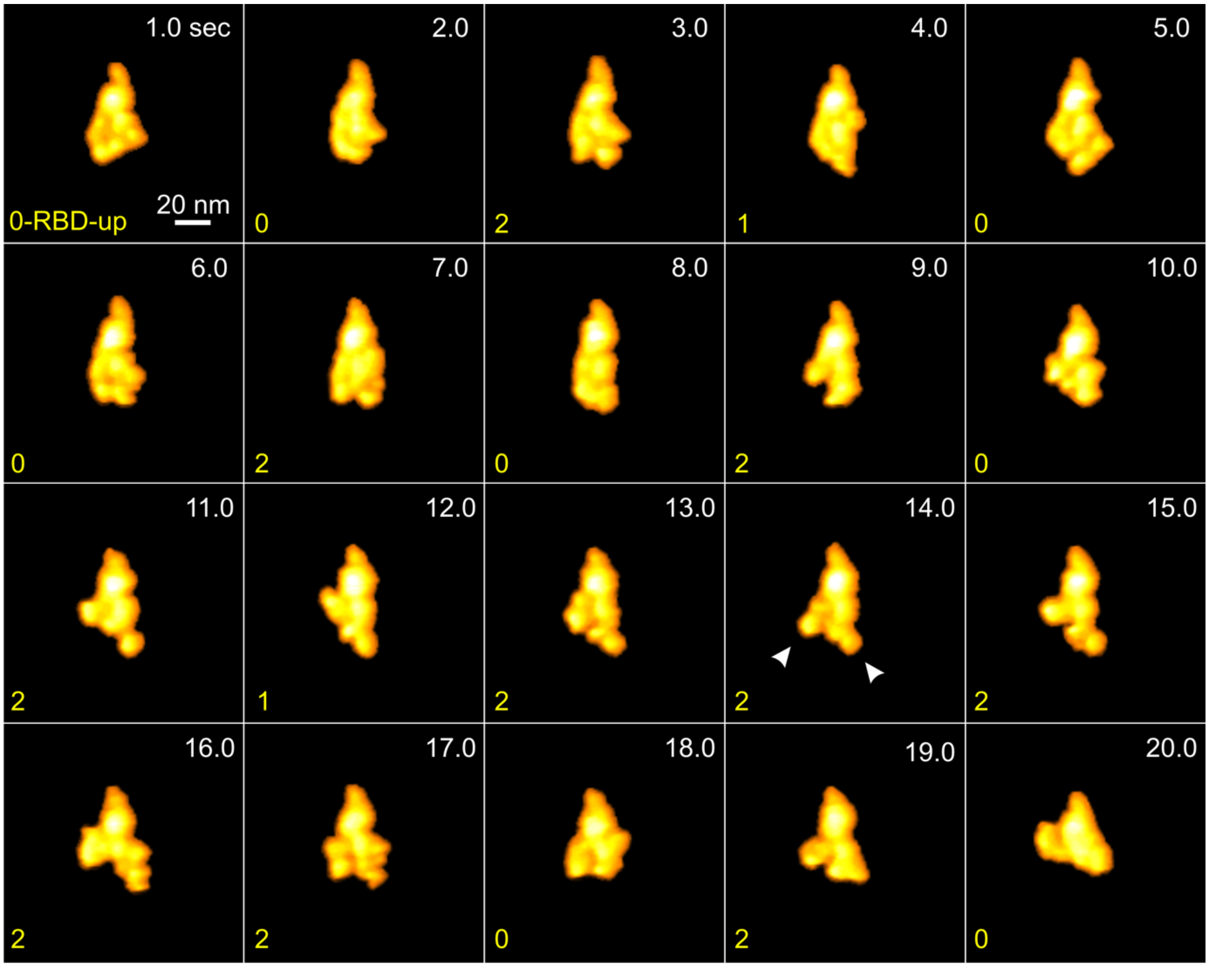


**Fig. S5.** Real-time $RBD$ conformations in $S^{f-6P}_{Omicron}$. Consecutive images from an HS-AFM video depicting $S^{f-6P}_{Omicron}$ in various states (as indicated in the bottom left corner of the images). The opening of $RBD$s is indicated with white arrowheads in one of the images at 14.0 s. The acquisition rate of the video was 1.0 fps. These images were processed with a set of user-written macros (see Materials and Methods) to reduce noise and remove small particles present in the background. Compared to the $S^{f-}_{Wuhan}$, the domains of the $S1$ subunit of $S^{f-6P}_{Omicron}$ appear to be more dynamic in nature (*41*).

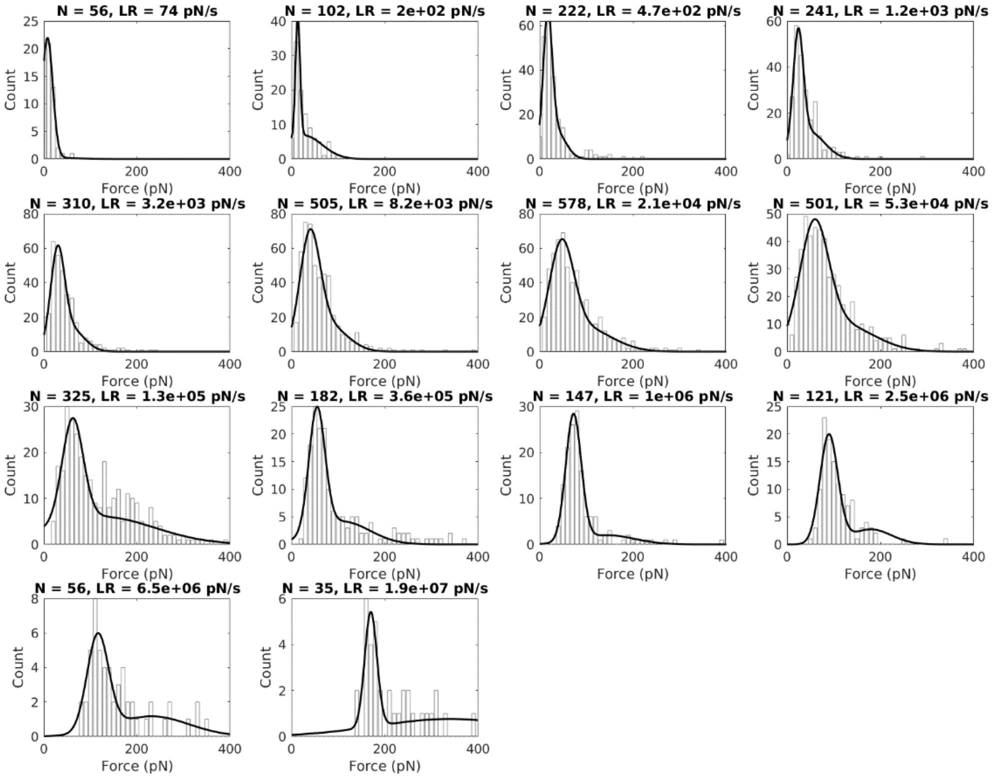


**Fig. S6.** Distributions of the rupture forces observed by HS-FS on individual complexes formed between $ACE2$ and $RBD_{Wuhan}$. Solid lines are best fits to double Gaussian distribution. The most probable force was determined as the maximum of the first peak and used to plot the dynamic force spectra shown in Fig. 3C. N is the number of events and LR is the loading rate.

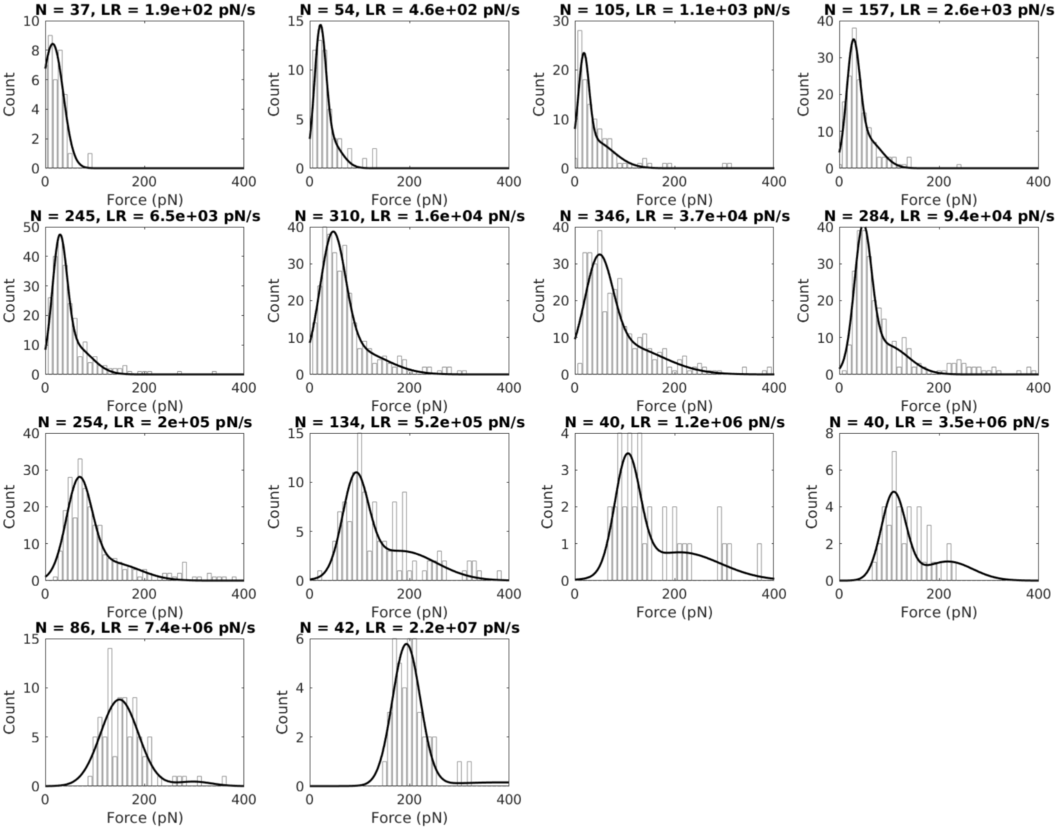


**Fig. S7.** Distributions of the rupture forces observed by HS-FS on individual complexes formed between $ACE2$ and $S_{Wuhan}^{f-}$. Solid lines are best fits to double Gaussian distribution. The most probable force was determined as the maximum of the first peak and used to plot the dynamic force spectra shown in Fig. 3C. N is the number of events and LR is the loading rate.

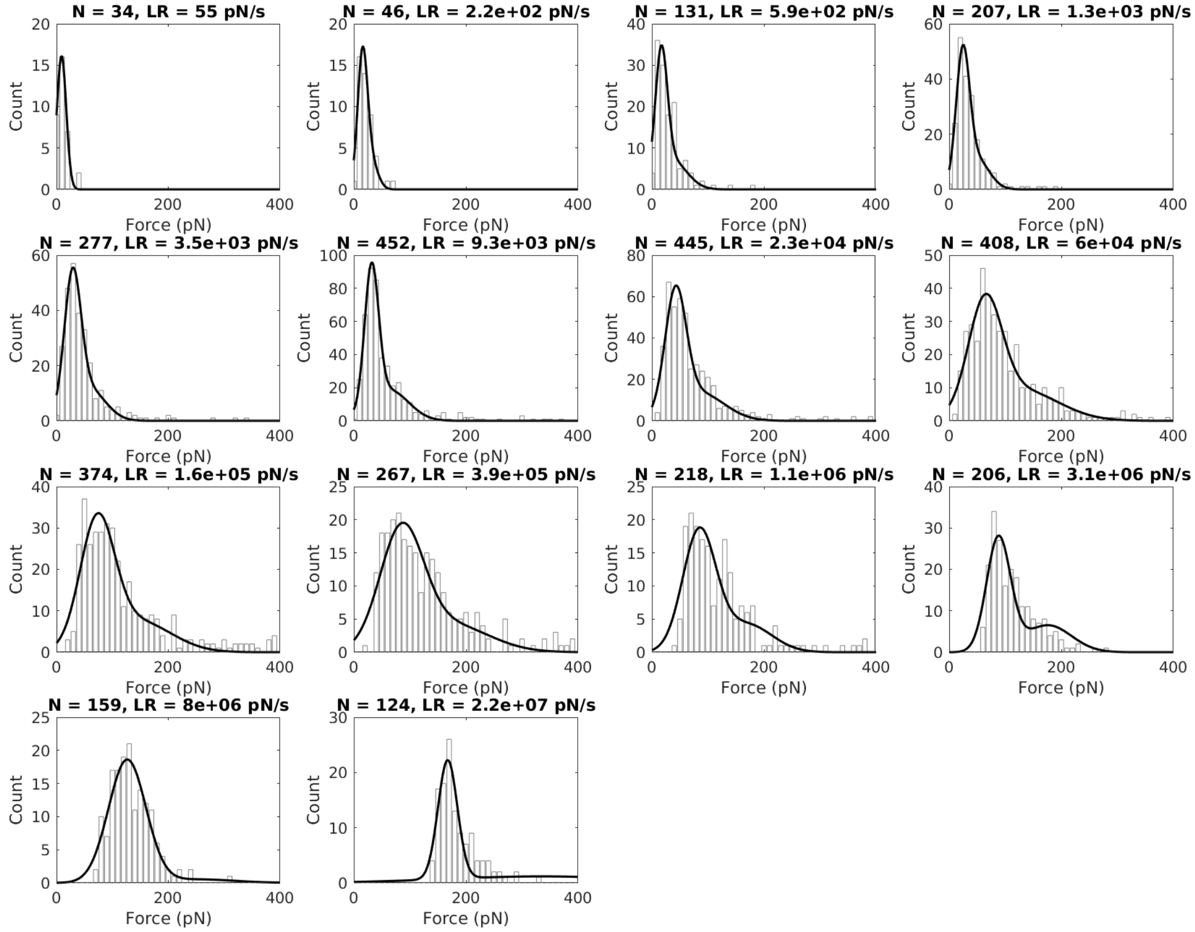


**Fig. S8.** Distributions of the rupture forces observed by HS-FS on individual complexes formed between $ACE2$ and $S_{Wuhan}^{2P}$. Solid lines are best fits to double Gaussian distribution. The most probable force was determined as the maximum of the first peak and used to plot the dynamic force spectra shown in Fig. 3C. N is the number of events and LR is the loading rate.

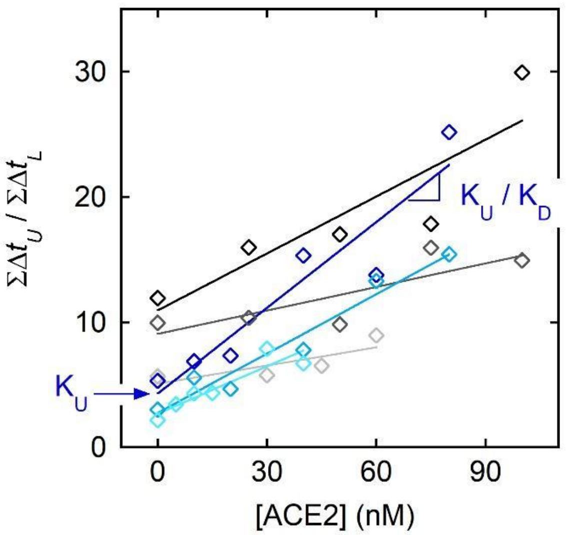


**Fig. S9.** Original experimental points and fits providing, from MT measurements at constant force, the dissociation equilibrium constant for the complexation reaction between the $ACE2$ and $RBD_{Wuhan}$, either isolated or embedded in $S$ trimer. The variations of the ratio between the total time spent in the unlooped conformation and the total time spent in the looped one were plotted as a function of the concentration in titrant. Data for three exemplary scaffolds where $ACE2$ faces $RBD_{Wuhan}$ were fitted according to Eq. Tit1, which, thanks to Eq. Tit2, results in the following dissociation equilibrium constant: $K_D$ = 17.5 ± 1.1, 18.8 ± 1.9, and 21.5 ± 2.7 nM (blue to cyan). Contrary to what can be seen for the same data in Fig. 4H, no renormalization had yet been performed here. Therefore, as indicated, a graphical reading of both $K_D$ and $K_U = K_D/C'_{eff}$ is possible. For the three other exemplary scaffolds where $ACE2$ faces $S^{2P}_{Wuhan}$ we obtained $K_D^{app}$ = 72.3 ± 10.9, 104.8 ± 24.1, and 145.3 ± 28.7 nM (black to grey).

**Table S1.** Populations of the different $RBD$ conformations of $S$ trimers from cryo-EM literature and from HS-AFM (this work). $K_C$ was calculated from the ratio of total fraction of $S$ protomers observed with all $RBD$s in the closed state ($\Sigma RBD_{closed} = \sum_i (3-i) \times S_i$) over those showing at least one $RBD$ in the open state ($\Sigma RBD_{open} = \sum_i i \times S_i$), $K_c \equiv \Sigma RBD_{closed} / \Sigma RBD_{open} = \sum_i (3-i) \times S_i / \sum_i i \times S_i$.

| **Variant** | **0-$RBD$ open** | **1-$RBD$ open** | **2-$RBD$ open** | **3-$RBD$ open** | $K_C$ | $p_{open}$ | **Ref.** |
|---|---|---|---|---|---|---|---|
| Wuhan | 66 | 34 | 0 | 0 | 7.8 | 0.11 | (*53*) |
| Wuhan 2P | 54 | 46 | 0 | 0 | 5.5 | 0.15 | (*54*) |
| Wuhan 2P | 54 | 46 | 0 | 0 | 5.5 | 0.15 | (*55*) |
| Wuhan 2P | 94 | 6 | 0 | 0 | 49.0 | 0.02 | (*56*) |
| Wuhan 2P | 45 | 55 | 0 | 0 | 4.5 | 0.18 | (*57*) |
| Wuhan 2P | 43 | 46 | 11 | 0 | 3.4 | 0.23 | (*58*) |
| Wuhan 2P + ACE2 | 0 | 42 | 53 | 5 | 0.9 | 0.54 | (*58*) |
| Wuhan 2P, -f | 51 | 49 | 0 | 0 | 5.1 | 0.16 | (*59*) |
| Wuhan D614G 2P | 5 | 36 | 39 | 20 | 0.7 | 0.58 | (*54*) |
| Wuhan D614G 2P | 18 | 82 | 0 | 0 | 2.7 | 0.27 | (*55*) |
| Wuhan D614G 2P | 5 | 36 | 29 | 20 | 0.8 | 0.57 | (*57*) |
| Wuhan D985C, S383C | 100 | 0 | 0 | 0 | ∞ | 0 | (*59*) |
| Wuhan G669C, T866C | 68 | 32 | 0 | 0 | 8.4 | 0.11 | (*59*) |
| Wuhan 2C | 100 | 0 | 0 | 0 | ∞ | 0 | (*60*) |
| Wuhan -f | 34 | 52 | 10 | 4 | 2.6 | 0.28 | Saha et al. (this work) |
| Wuhan 2P | 39 | 54 | 6 | 1 | 3.3 | 0.23 | Saha et al. (this work) |
| Wuhan 2C | 76 | 22 | 2 | 0 | 10.5 | 0.09 | Saha et al. (this work) |
| Wuhan 2P, 2C | 59 | 36 | 5 | 0 | 5.5 | 0.15 | Saha et al. (this work) |
| Omicron | 40 | 50 | 0 | 0 | 4.4 | 0.19 | (*46*) |
| Omicron -f | 42 | 35 | 23 | 0 | 2.7 | 0.27 | (*61*) |
| Omicron 2P | 100 | 0 | 0 | 0 | ∞ | 0 | (*41*) |
| Omicron 2P | 0 | 100 | 0 | 0 | 2.0 | 0.33 | (*57*) |
| Omicron 2P | 0 | 45 | 55 | 0 | 0.9 | 0.52 | (*62*) |
| Omicron 6P | 1 | 74 | 12 | 12 | 1.2 | 0.45 | (*63*) |
| Omicron 6P | 54 | 32 | 12 | 1 | 4.0 | 0.20 | Saha et al. (this work) |

**Table S2.** Kinetic and thermodynamic parameters for the closing transition of an individual $RBD$ within the $S$ trimer. The closing equilibrium constant, $K_C$, was calculated from three different approaches. First, from the ratio of the total number of images showing $S$ trimer with all $RBD$s in the closed state ($\Sigma RBD_{closed} = \sum_i (3-i) \times S_i$) over those showing at least one $RBD$ in the open state ($\Sigma RBD_{open} = \sum_i i \times S_i$). Second, from the probability for $RBD$ to be in an open state, as obtained from the fit of the binomial distribution $S_i = \frac{3!}{i!(3-i)!} \times p_{open}{}^i \times (1-p_{open})^{3-i}$ to the relative occupancy histograms (Fig. 1C). Third, from the ratio between the closing and opening rates extracted from the global fits to the dwell time distributions (Fig. 1E).

| $S$ Variant | $K_c \equiv \frac{\Sigma RBD_{closed}}{\Sigma RBD_{open}}$ | $p_{open}$ | $K_C \equiv \frac{1}{p_{open}} - 1$ | $k_C$ ($s^{-1}$) | $k_O$ ($s^{-1}$) | $K_C \equiv \frac{k_C}{k_O}$ | N videos | N images | N transitions |
|---|---|---|---|---|---|---|---|---|---|
| $S^{f-}_{Wuhan}$ | 2.6 ± 0.6 | 0.28 ± 0.04 | 2.5 ± 0.4 | 1.00 ± 0.28 | 0.40 ± 0.11 | 2.5 ± 1.4 | 13 | 2780 | 930 |
| $S^{2P}_{Wuhan}$ | 3.3 ± 0.8 | 0.26 ± 0.04 | 2.9 ± 0.6 | 0.88 ± 0.36 | 0.52 ± 0.21 | 1.7 ± 1.4 | 6 | 816 | 230 |
| $S^{f-,2C}_{Wuhan}$ | 10.5 ± 2.0 | 0.09 ± 0.00 | 10.4 ± 0.0 | 1.81 ± 0.60 | 0.44 ± 0.15 | 4.2 ± 2.8 | 9 | 1357 | 461 |
| $S^{f-,2C,2P}_{Wuhan}$ | 5.5 ± 1.9 | 0.16 ± 0.01 | 5.1 ± 0.2 | 1.04 ± 0.39 | 0.38 ± 0.14 | 2.7 ± 2.0 | 7 | 580 | 189 |
| $S^{f-,6P}_{Omicron}$ | 4.0 ± 0.6 | 0.18 ± 0.01 | 4.6 ± 0.4 | 1.12 ± 0.26 | 0.21 ± 0.05 | 5.2 ± 2.5 | 18 | 1361 | 572 |

**Table S3.** Energy landscape parameters for the dissociation reaction between the $ACE2$ and $RBD$, either isolated or embedded within the $S$ trimer. $k_D$ corresponds to the dissociation rate constants, $x^‡$ to the distance to the transition state, $\Delta G^‡$ and $E_D$ to the free and activation energies, respectively. HS-FS experiments were performed at room temperature in PBS at pH 7.4 and MT experiments, at 30 °C in 10 mM Na-HEPES pH 7.4, 150 mM NaCl, 0.1 % Tween 20, and 0.5 mg/mL BSA.

| **Setup** | **Viral partner** | $k_D$ **($10^{-3}$ s$^{-1}$)** | $x^‡$ **(nm)** | $\Delta G^‡$ **or** $E_D$ **($k_BT$)** |
|---|---|---|---|---|
| HS-FS | $RBD_{Wuhan}$ | 6 ± 2 | 1.06 ± 0.09 | 15.3 ± 1 |
| | $S^{f-}_{Wuhan}$ | 11 ± 6 | 0.84 ± 0.08 | 14.9 ± 1 |
| | $S^{2P}_{Wuhan}$ | 4 ± 2 | 1.12 ± 0.07 | 16.1 ± 1 |
| MT | $RBD_{Wuhan}$ | 23 ± 2 | 0.66 ± 0.07 | 54.1 ± 8 |
| | $S^{2P}_{Wuhan}$ | 22 ± 2 | 0.79 ± 0.12 | 44.4 ± 6 |

**Supplementary videos (SV) information.**

List of HS-AFM videos that show $RBD$-based conformational dynamics of the $S$ trimer variants, as detailed in this study. Each video has a time stamp in the top-left corner and the x-y scale in the bottom-right corner. The presence of protrusion, a signature of $RBD$ opening, is denoted with a cyan contour in the videos and displayed alongside an unmarked one. In cases where the video depicts all $RBD$ in closed state, only the unmarked version is provided.

| Video | $S$ variant | Acquisition rate (fps) | Display rate (fps) |
|---|---|---|---|
| SV1 | $S_{Wuhan}^{f-}$ | 1 | 1 |
| SV2 | $S_{Wuhan}^{f-}$ | 2.5 | 5 |
| SV3 | $S_{Wuhan}^{f-}$ | 1 | 5 |
| SV4 | $S_{Wuhan}^{f-}$ | 1 | 5 |
| SV5 | $S_{Wuhan}^{f-}$ | 1 | 5 |
| SV6 | $S_{Wuhan}^{f-}$ | 0.5 | 5 |
| SV7 | $S_{Wuhan}^{2P}$ | 2 | 5 |
| SV8 | $S_{Wuhan}^{2P}$ | 1 | 1 |
| SV9 | $S_{Wuhan}^{2P}$ | 1 | 5 |
| SV10 | $S_{Wuhan}^{f-,2C}$ | 2.5 | 5 |
| SV11 | $S_{Wuhan}^{f-,2C}$ | 1 | 3 |
| SV12 | $S_{Omicron}^{f-,6P}$ | 1 | 5 |
| SV13 | $S_{Omicron}^{f-,6P}$ | 1 | 5 |
| SV14 | $S_{Omicron}^{f-,6P}$ | 1 | 5 |
| SV15 | $S_{Omicron}^{f-,6P}$ | 1 | 5 |
| SV16 | $S_{Omicron}^{f-,6P}$ | 1 | 5 |
| SV17 | $S_{Wuhan}^{f-,2C,2P}$ | 2.5 | 5 |
| SV18 | $S_{Wuhan}^{f-,2C,2P}$ | 1 | 5 |
| SV19 | $S_{Wuhan}^{f-,2C,2P}$ | 1 | 1 |
| SV20 | $S_{Wuhan}^{f-}$ global view | 1 | 5 |
